# Disassembly of hemidesmosomes promotes tumorigenesis in PTEN-negative prostate cancer by targeting plectin into focal adhesions

**DOI:** 10.1101/2021.11.08.467671

**Authors:** Tomasz Wenta, Anette Schmidt, Qin Zhang, Raman Devarajan, Prateek Singh, Xiayun Yang, Anne Ahtikoski, Markku Vaarala, Gong-Hong Wei, Aki Manninen

## Abstract

Loss of α6β4-dependent hemidesmosomes has been observed during prostate cancer progression. However, the significance and underlying mechanisms by which aberrant hemidesmosome assembly may modulate tumorigenesis remain elusive. Using an extensive CRISPR/Cas9-mediated genetic engineering approaches in different prostate cancer cell lines combined with *in vivo* tumorigenesis studies in mice, bone marrow-on-chip assays and bioinformatics, as well as histological analysis of prostate cancer patient cohorts, we demonstrated that simultaneous loss of PTEN and hemidesmosomes induced several tumorigenic properties including proliferation, migration, resistance to anoikis, apoptosis, and drug treatment *in vitro*, and increased metastatic capacity *in vivo*. Our studies showed that these effects were driven by activation of EGFR/PI3K/Akt and FAK/Src-pathways and were abolished by plectin downregulation. Therefore, dual loss of PTEN and hemidesmosomes may have diagnostic value helping to stratify prostate cancer patients with high risk for development of aggressive disease and highlight plectin as a potential therapeutic target in prostate cancer.

## Introduction

Prostate cancer (PCa) is the second most common cancer in men with more than a million new cases diagnosed annually ^1^. A comprehensive genetic landscape of PCa tumorigenesis and susceptibility has been pieced together ^2–5^, and these studies have recently been extended to investigations on the proteogenomic landscape ^6, 7^. Due to improved screening and awareness, many PCa are detected while still localized enabling effective surgical or radiotherapy-based treatments. However, these radical treatments are frequently associated with severe side effects that are a considerable concern because a significant fraction of diagnosed prostate tumors are indolent ^8, 9^. Therefore, a better understanding of the molecular signatures differing between indolent and aggressive tumors is urgently needed.

The gradually changing microenvironment surrounding cancer cells has emerged as a critical driver of tumorigenesis ^10^. PCa progression is linked with loss of hemidesmosomes (HD) ^11–13^. It is not known when or how the loss of HDs takes place during PCa pathogenesis. α6β4-integrins are the crux of HDs and earlier data suggested that loss of β4-integrin expression in PCa cells leads to the disintegration of HDs and release of α6-integrin allowing it to pair with β1-integrin, thereby driving invasive migration ^14^. However, it remains unclear whether, and in which context, loss of HDs is the cause (active role) or a consequence (passive indicator) of PCa progression.

Here we have addressed the role of HD disassembly in PCa by studying HDs in both benign and malignant prostate epithelial cells. The distribution pattern of HDs was altered in prostate carcinoma cells studied, yet HD components remained colocalized. Disruption of HDs by depletion of integrin α6- or β4-subunits led to redistribution of HD components. Interestingly, expression of plectin, an HD protein responsible for linking HDs to the intermediate filament (IF) network, was decreased in benign but increased in malignant prostate epithelial cells upon loss of HDs. We found that plectin upregulation in HD-depleted cells occurred specifically in the absence of PTEN expression, a well-known tumor suppressor frequently lost in PCa. In the absence of HDs and PTEN, plectin was retargeted towards actin-linked focal adhesions (FA) and was found to induce FA-signaling, cell migration, and proliferation. Loss of HDs in PTEN-negative prostate carcinoma cells also promoted drug- and anoikis-resistance and enhanced their tumorigenic potential both *in vitro* and *in vivo*. Importantly, we showed that simultaneous loss of HDs and PTEN is sufficient to transform benign prostate cells promoting their capability for metastasis in mice *in vivo*. Finally, our findings were corroborated in a clinical setting by an extensive analysis of independent cohorts with PCa and a tissue microarray analysis of an additional PCa patient cohort to show that loss of PTEN and HD assembly significantly correlates, in plectin-dependent manner, with an aggressive form of PCa and worse overall survival of PCa patients.

## Results

### Hemidesmosome organization is altered in malignant PCa cells

First, we assessed α6β4-integrin expression in normal and PCa epithelial cell lines. In line with previous reports ^13, 15^, benign cells (RWPE1) expressed the highest level of hemidesmosomal (HD) α6- and β4-integrins whereas PCa cells had drastically reduced expression of integrin α6- and especially β4-subunit (Figure 1A). To compare HD organization in benign and malignant prostate epithelium we selected normal (RWPE1) and malignant (PC3 and DU145) prostate epithelial cells, all of which express α6β4-integrins, and stained them for selected HD components. RWPE1 is an immortalized cell line from the normal prostate whereas PC3 (grade IV bone metastasis) and DU145 (grade II brain metastasis) are adenocarcinoma cell lines derived from metastatic lesions. In confluent RWPE1 cells, α6β4-integrin antibodies showed a nearly uniform basal staining that was interrupted by few circular “holes” (Figure 1B). These α6β4-integrin negative areas had prominent actin staining and thus likely represent podosomes (van den Dries et al., 2013). Known HD components, plectin and CD151, colocalized with α6β4-integrins at HDs (Figures 1C, 2B) whereas focal adhesion (FA) components, forming another integrin-based adhesion complex, localized mainly to lateral peripheral borders of confluent RWPE1 cells with essentially no overlap with HDs, confirming that these are distinct structures (Figure 1C).

**Figure. 1.**
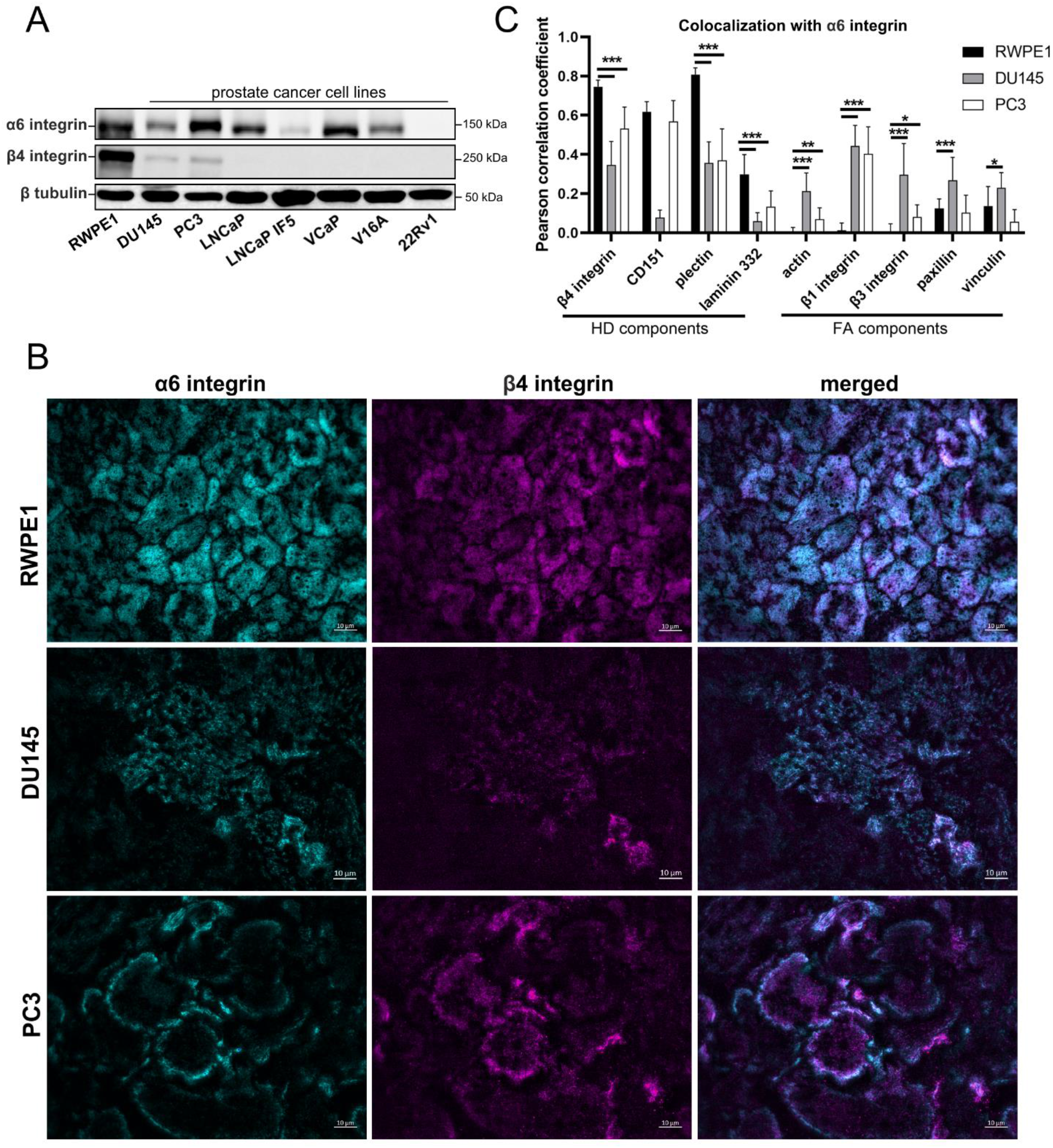
HDs organization is altered in PCa cell lines. Level of HD-associated α6β4-integrin in normal (RWPE1) and PCa (DU145, PC3, LNCap, LNCaP-IF5, VCaP, V16A, 22Rv1) epithelial cells (A); Immunofluorescence analysis of the subcellular localization of integrin α6- (cyan) and β4-subunits (magenta) in normal (RWPE1) and PCa (DU145, PC3) cells (B); Pearson correlation coefficient (PCC) analysis to measure the colocalization of α6-integrin with the indicated HD or FA components (C). The data are presented as mean ± SD, n≥20.

**Figure 2.**
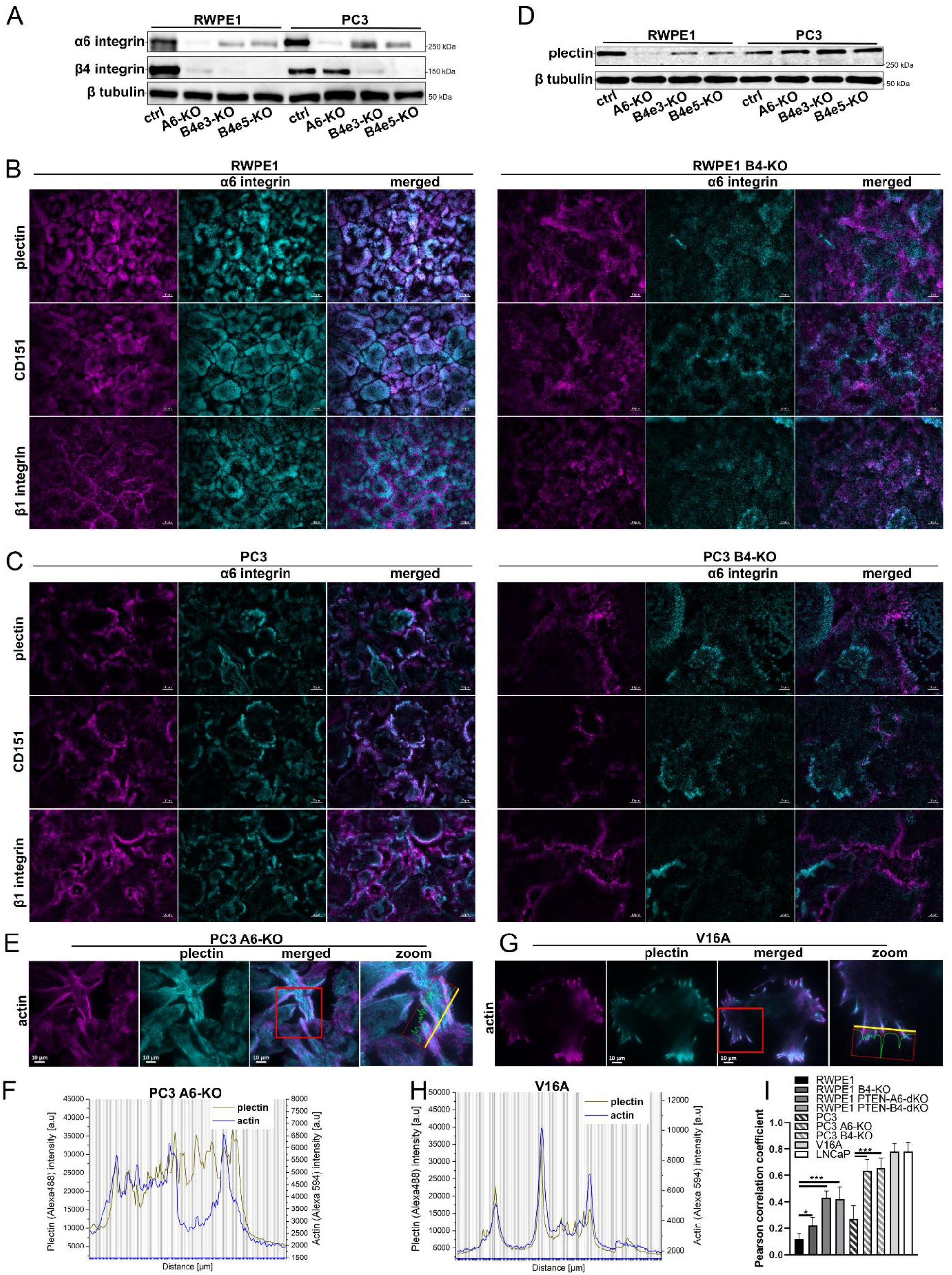
The outcome of integrin α6- or β4-subunit depletion in normal and cancer prostate epithelial cells. α6- and β4-integrin subunits are stabilized by heterodimerization in normal (RWPE1) but to a lesser extent in cancer (PC3) cells (A); Immunofluorescence analysis of α6-integrin in RWPE1-ctrl and RWPE1 β4-KO cells (B); Immunofluorescence analysis of α6 integrin in PC3 and PC3 β4-KO cells (C); Level of plectin in α6- or β4-integrin depleted RWPE1 and PC3 cells (D); Immunofluorescence analysis of colocalization of plectin with actin in HD-deficient PC3 α6-KO (E) and V16A (G); Plot of plectin-actin colocalization in PC3 α6-KO (F) and V16A cells (H); Comparison of plectin-actin Pearson correlation coefficient value in HD-deficient cells (I). The data are presented as mean ± SD, n≥20.

Although α6β4-integrins were still expressed in malignant prostate cells and colocalized at least partially with other HD components, the HD organization was clearly different (Figures 1B-C). In PC3 cells, HDs formed peripheral rings while in DU145 cells HD organization was more similar to RWPE1 but appeared more fragmented (Figure 1B). In DU145 cells, α6/CD151 colocalization and the surface level of β4-integrin detected using TIRF-microscopy was significantly lower than in the other two cell lines studied. Curiously, two distinct patterns of α6-integrin staining were observed in DU145 cells. Most cells displayed α6-staining colocalizing with β4-integrins, but in some cells α6-integrin localized to small intense spots with an FA-like organization (Figure 1B). In PC3 cells, α6β4-integrins and CD151 colocalized at peripheral adhesive domains whereas plectin showed a moderate reduction in colocalization with HDs extending slightly more peripherally (Figures 1B-C, 2C).

### Disruption of HDs by α6- or β4-integrin depletion leads to alterations of plectin level and its distribution in prostate epithelial cells

To investigate the functional role of HDs in prostate epithelial cells we disrupted them by knocking-out α6 or β4-integrin subunit using CRISPR/Cas9-mediated genome editing (Figure 2A). Deletion of β4-integrin in RWPE1 cells led to a drastic reduction at α6-integrin expression level (Figure 2A). Similarly, β4-integrin expression levels were strongly downregulated in α6-KO RWPE1 cells. These data suggest that the formation of α6β4-integrin heterodimer is needed for the stability of individual α6- and β4-chains in normal prostate epithelial cells. Loss of HDs in RWPE1 led also to loss of podosomes and downregulation of plectin (Figures 2B, 2D, S1). Another HD component CD151 was redistributed to cell-cell junctions (Figures 2B, S1).

Interestingly, loss of either α6- or β4-integrins in PC3 cells did not result in equally strong downregulation of their heterodimer partner (Figure 2A). Loss of β4-integrin in PC3 resulted in reduced and diffuse basal staining of α6-integrin (Figure 2C). In α6-integrin knock-out (α6-KO) cells β4-integrin was largely excluded from the basal membrane (Figure S1). Surprisingly, plectin expression was increased in both α6- and β4-depleted PC3 cells and it was found distributed at the basal membrane in a stripy FA-like pattern (Figures 2C-D, S1). Since plectin has been shown to interact also with the actin cytoskeleton ^16^, we stained actin in these cells. The immunofluorescence analysis of PC3 α6- or β4-KO showed an increased abundance of basal actin stress fibers (Figure 2E) linked to focal adhesions (FA). Plectin localized close to FAs in PC3 cells but it did not completely colocalize with FA markers or actin (Figures 2E-F, 2I). We made similar observations in HDs-deficient V16A (β4-integrin-negative) cells derived from the LNCaP cancer cell line ^17^. In HD-disrupted V16A cells plectin colocalized partially with actin fibers (Figures 2G-I). Overall, comparison of the Pearson correlation coefficients for plectin-actin colocalization, revealed significantly higher colocalization in cells deficient for HD assembly (Figure 2I).

Collectively, this analysis showed that malignant PCa cells (PC3, DU145) do form HD-like structures that are differently organized when compared with HDs in benign RWPE1 cells. Depletion of either α6- or β4-integrin led to alteration of the other and of plectin. In control cells, plectin levels were downregulated upon HD disruption whereas in malignant cells plectin was upregulated and redistributed towards FAs mainly at the basal cell-cell borders.

### Loss of HDs activates FA- and EGFR-signaling in PTEN-negative cancer cells

Given the observed proximity of plectin and FAs in HD-depleted cells, we next analyzed PC3 and PC3 α6-KO, and PC3 β4-KO for possible changes in the expression levels of FA proteins and in FA-mediated signaling. The protein levels of vinculin and paxillin levels were upregulated in HD-depleted PC3 cells (Figure 3A). The levels of active phosphorylated Src (pSrc^Y4^^16^) and phosphorylated focal adhesion kinases (pFAK^Y397^) were also upregulated, suggesting activation of integrin-mediated signaling (Figure 3A). Moreover, the levels of paxillin, a key FA protein, were drastically upregulated (Figure 3A). Upregulation of vimentin expression was also noted in HD-depleted PC3 cells. One of the downstream targets of integrin activation is Akt. A strong activation of Akt was detected in PC3 α6-KO cells while PC3 β4-KO cells showed a more modest activation (Figure 3A). Moreover, in both α6- and β4KO cells we observed robust activation of epidermal growth factor receptor (EGFR) that has been reported to drive PI3K-Akt phosphorylation leading to MAPK signaling activation in synergy with integrin ligation (Gu et al., 1998). Both EGFR and MAPK-signaling were upregulated in HD-depleted PC3 cells. EGFR-signaling and its downstream PI3K-Akt pathway are counteracted by PTEN, a tumor suppressor frequently lost in the most aggressive PCa cells ^18^, including PC3 cells that are derived from a PTEN-negative PCa bone metastasis ^19^. Thus, our data suggest that in the absence of functional PTEN (in PC3), HDs can limit PI3K-Akt activation, and the disassembly of HDs is followed by activation of EGFR/PI3K/Akt/MAPK-signaling, presumably via stimulation of FA-associated signaling.

**Figure 3.**
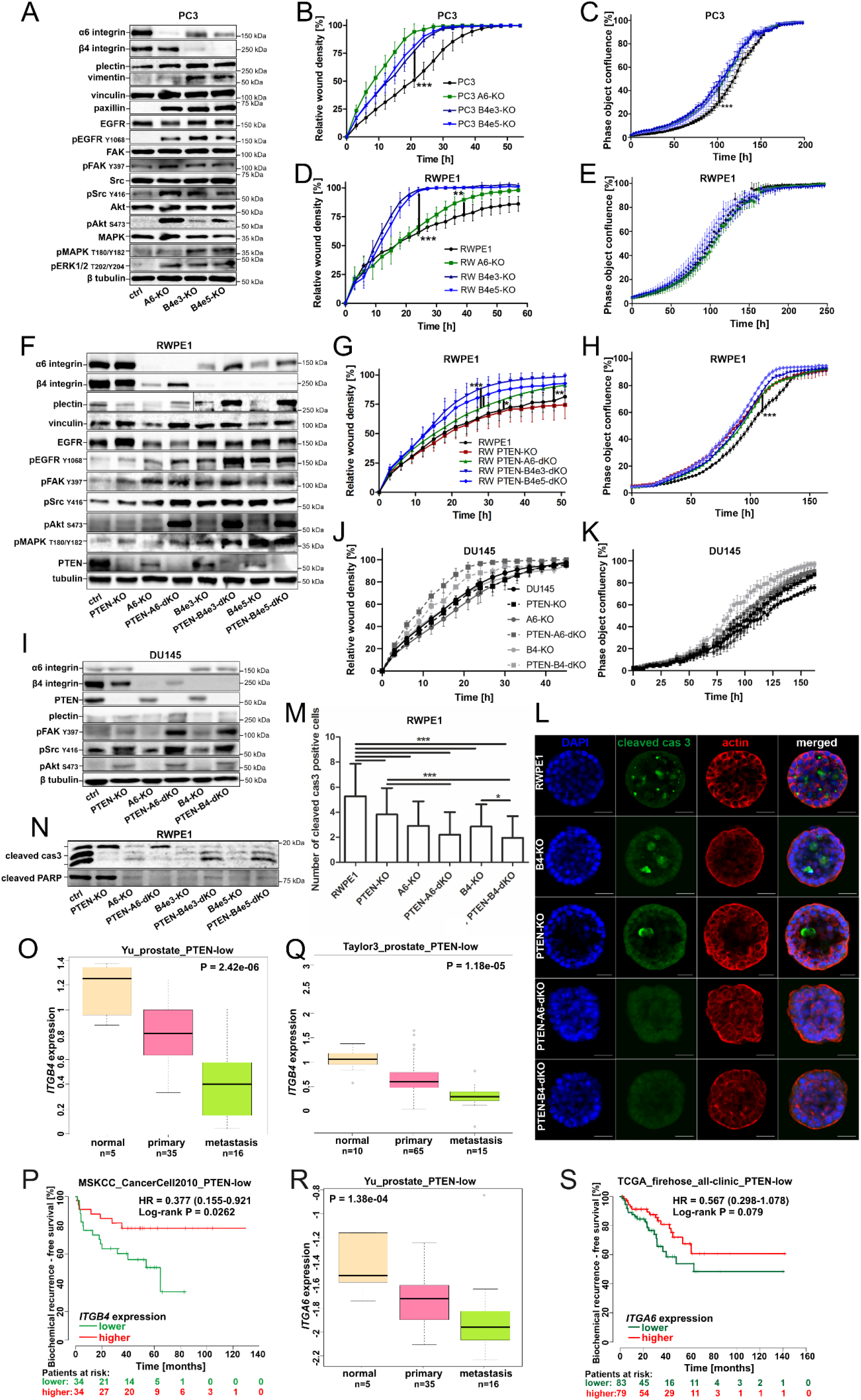
Loss of HDs promotes cell migration, stimulates FA-mediated integrin signaling and cell proliferation in PTEN-negative cancer cells. Western blotting analysis of PC3 (A), RWPE1 and RWPE1 PTEN-KO (F), and DU145 and DU145 PTEN-KO (I) cell lines with α6- or β4-KO; The effect of HD disruption on wound healing ability in PC3 (B) RWPE1 (D), RWPE1 PTEN-KO (G) and DU145 (J) cell lines; the effect of α6- or β4-KO on proliferation in PC3 (C) RWPE1 (E), RWPE1 PTEN-KO (H) and DU145 (K) cell lines; 3D culture of parental and RWPE1 mutant cell lines (L); Quantitative analysis of apoptosis (luminal cells positive for cleaved caspase-3 staining) in 3D cultures (M); western blot analysis of apoptotic markers of the indicated cell lines grown on Poly-HEMA-coated plates (N); ITGB4 (O-Q) or ITGA6 (R-S) is downregulated upon PCa progression and associated with poorer prognosis in stratified PTEN-low PCa patients. The data are presented as mean ± SD. All analyses were performed in triplicate.

To study the effect of increased FA- and EGFR/Akt-signaling in HD-depleted PC3 cells, we analyzed the migratory and proliferative capacity of the different PC3 cell lines. We observed that both PC3 α6-KO and β4-KO cells migrated and proliferated faster than parental PC3 cells suggesting increased tumorigenic potential (Figures 3B-C). To study if this effect is limited to malignant cells, we analyzed migration and proliferation of α6- or β4-integrin-depleted benign RWPE1 cells and found that while migration was induced, particularly in β4-deficient cells (Figure 3D), there was no effect on proliferation (Figure 3E). Therefore, the full spectrum of enhanced tumorigenic potential upon HD-depletion was only seen in malignant tumor cells (Figures 3B-C). As mentioned previously, one of the key genetic differences between RWPE1 and PC3 cells is their PTEN-status ^19^. To investigate the potential role of PTEN-status, we deleted PTEN in RWPE1, RWPE1 α6-KO and RWPE1 β4-KO cells (Figure 3F). Deletion of PTEN from RWPE1 cells stimulated their proliferation but did not affect migration (Figures 3G-H). In contrast, dual loss of PTEN and α6- or β4-integrin not only induced proliferation but also promoted FA-signaling and cell migration (Figures 3F-H). Moreover, co-deletion of PTEN and α6- or β4-integrin in DU145 cells, a PCa cell line that is heterozygous for functional PTEN, phenocopied PC3 α6-KO and β4-KO cells confirming that HD disruption in PTEN-negative cells induces FA-mediated signaling, cell proliferation and migration (Figures 3I-K). Consistent with this model, re-expression of α6-integrin in RWPE1-PTEN-α6-dKO or PC3 α6-KO restored their migratory capacity down to similar levels with their respective parental control cells, while overexpression of α6-integrin in PC3 cells had no significant effect (Figure S2).

Finally, to study if simultaneous loss of PTEN and HDs may induce FA signaling also in other cancer types, we introduced the mutations to JIMT-1 breast cancer cells with intact PTEN-function. In agreement with the data from PCa cells, dual loss of PTEN and HDs in JIMT-1 cells induced FA-signaling, Akt and prevented the downregulation of plectin levels observed in HD-depleted (PTEN-positive) JIMT-1 cells (Figure S3).

### PTEN-loss and HD-disassembly synergistically promote anoikis-resistance and progression of an aggressive PCa

Three-dimensional (3D) culture system can be used to study subtle defects in epithelial cell polarity as well as anoikis-resistance of cells shed into the apical lumen ^20^. RWPE1 cells efficiently form round hollow cysts with relatively frequent apoptotic cells in the lumen stained positive for cleaved caspase 3 (Figure 3L). RWPE1 α6- and β4-KO cells with disrupted HDs, and PTEN-KO cells were all found to have significantly fewer apoptotic cells (Figures 3L-N). However, cells with a combined loss of HDs and PTEN were the most resistant to apoptosis in this model (Figures 3L-N). We confirmed these results with an independent method by culturing cells on non-adherent conditions (Poly-HEMA assay) followed by detection of cleaved caspase 3 and PARP staining in the cell lysates. This analysis showed that caspase 3 activation was inhibited whereas PARP-cascade activation was unaffected in PTEN-KO cells (Figure 3N). Curiously, HD disruption abrogated both pathways (Figure 3N). To further explore if these findings indicate any clinical relevance, we stratified several independent PCa cohorts in accordance with their PTEN-status. The data stratification was performed regarding PTEN expression level (Figures 3O-S, S4) and PTEN copy number loss (Figure S5). In line with our experimental data, bioinformatic analysis revealed that *ITGA6* and *ITGB4* expression levels negatively correlated with both metastatic disease and worse prognosis for biochemical recurrence indicating a progressive disease in low PTEN level (Figures 3O-S, S4) groups of PCa patient’s tissues with low levels of either *ITGA6* or *ITGB4*. Importantly, in patients with PTEN copy loss reduction of *ITGB4* or *ITGA6* expression significantly increased PCa aggressiveness (Figure S5).

Taken together, our data show that dual loss of HD assembly and PTEN promotes several tumorigenic properties by inducing cell migration, proliferation, and anoikis-resistance and suggest that disruption of HDs is particularly detrimental in the context of inactive PTEN function, a condition that is one of the most common genomic aberrations in PCa.

### Loss of HDs enhances tumorigenic potential of PTEN-negative cells in vitro and in vivo in a plectin-dependent manner

Since our data revealed an intriguing upregulation and FA-proximal targeting of plectin (Figure 2), we next investigated the potential functional role of plectin in mediating the effects of dual HD/PTEN-loss by introducing plectin-KO into PC3 α6-KO and β4-KO cells. Depletion of plectin (PLEC-KO) efficiently prevented upregulation of FA-mediated signaling (Figure 4A), cell proliferation (Figure 4B) and cell migration (Figure 4C) in PC3 α6-KO and β4-KO cells, suggesting that the plectin expression was indeed required for these effects. The critical role of plectin in HD-deficient cells on migration was also confirmed by deleting plectin in highly migratory RWPE1 PTEN-α6-dKO and RWPE1 PTEN-β4-dKO cells to generate triple KO cells with significantly inhibited migration even when compared with parental RWPE1 cells (Figure 4D).

**Figure 4.**
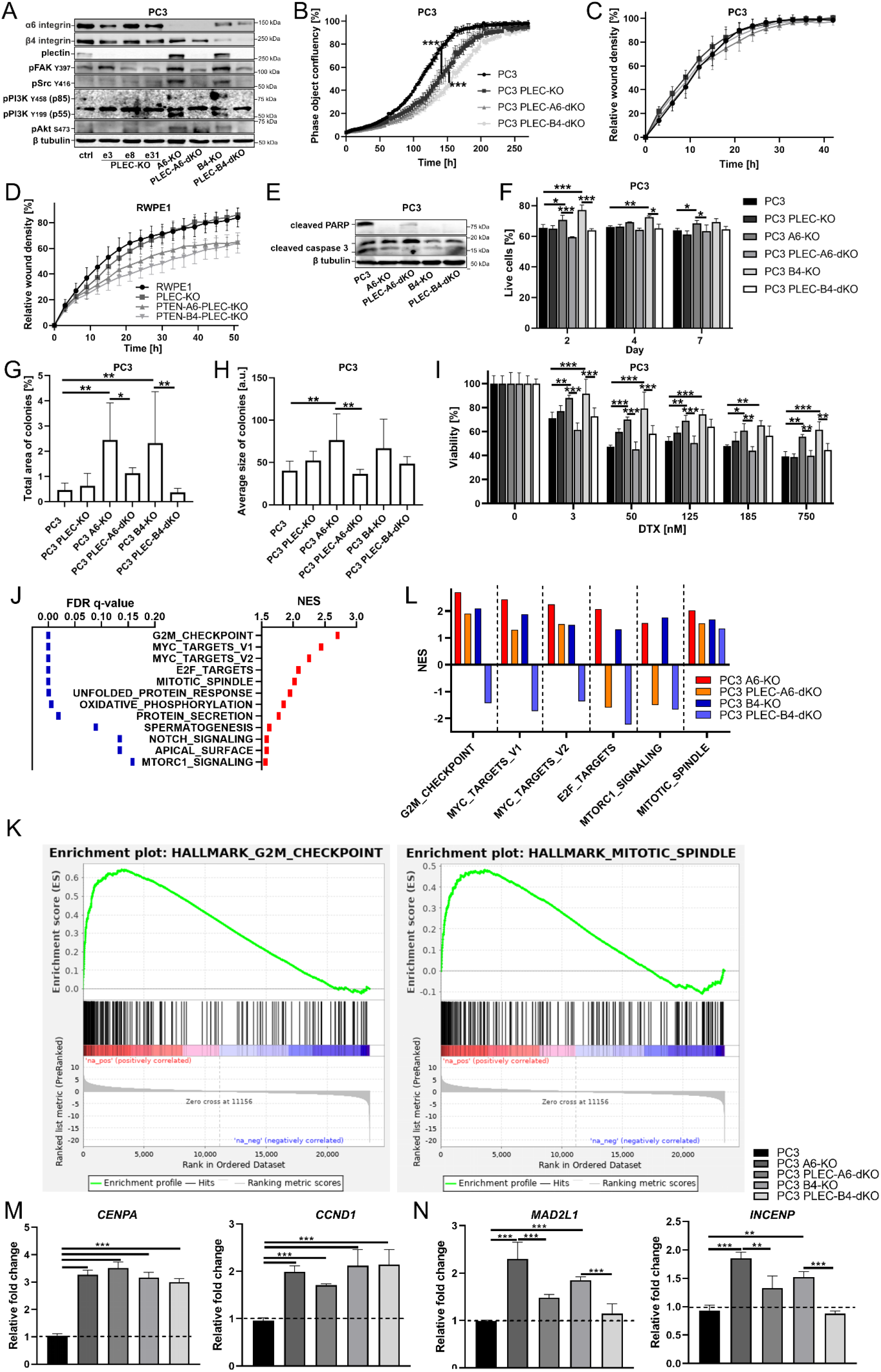
Plectin is required for the tumorigenic properties induced by dual loss of HDs and PTEN. Western blot analysis of selected signaling proteins in PC3, PC3-PLECKO, PC3-α6KO, PC3-PLEC-α6dKO, PC3-β4KO and PC3-PLEC-β4dKO cell lines (A); The effect of PLECKO on the wound healing ability (B) and proliferation (C) of the indicated PC3 cell variants; wound healing analysis of RWPE1, RWPE1-PTEN-α6-PLEC-tKO and RWPE1-PTEN-β4-PLEC-tKO cells (D). Analysis of apoptotic markers (E) anoikis-resistance (F-H) and docetaxel-resistance (I) in the indicated PC3 cell variants. Analysis of anoikis-resistance of the indicated PC3 cell line variants using PolyHEMA coated plates (F) Colony-forming soft agar assay with analysis of the total area of colonies (G) and an average size of individual colonies (H). Top enriched pathways upregulated in HD-deficient PC3-α6KO cells (J); Enrichment plots of the 1^st^- and 5^th^-ranked pathways in PC3 α6KO cells (K); Comparison of enriched pathways in the indicated PC3 cell variants (L); Quantitative PCR analysis confirming upregulation of selected markers of the G2M checkpoint pathway in plectin-independent manner (M) and in plectin-dependent manner (N). The data are presented as mean ± SD. All analyses were performed in triplicate.

To further assess the tumorigenic potential and plectin-dependence of HD-depleted PC3 cells we studied their anoikis-resistance by utilizing two independent methods, the poly-HEMA assay and the soft agar colony-formation assay. Despite being cancer cells, PC3 cells also activate apoptotic pathways when cultured on poly-HEMA-coated plates as shown by induction of cleaved caspase 3 and cleaved PARP levels under these conditions (Figure 4E). Both PC3 α6-KO and β4-KO cells displayed strong resistance to PARP-cleavage that appeared partially plectin-dependent, whereas the effect on caspase 3 cleavage was somewhat weaker (Figure 4E). These results were reflected by the increased percentage of live HD-depleted cells, when compared with the controls, in the flow cytometric analysis of cells harvested from the poly-HEMA-coated plates (Figure 4F). Importantly, this functional analysis indicated that HD-disassembly-mediated effect on apoptosis resistance was abolished in PLEC-KO cells. Analysis of soft agar-grown colonies also confirmed a significant plectin-dependent effect of HD-depletion on anoikis-resistance, as PC3 α6-KO and β4-KO cells formed more and significantly larger colonies when compared with parental PC3 cells (Figures 4G-H). Next, we extended these studies to assessment of drug resistance of PC3, α6-KO and β4-KO cells by exposing them to different concentrations of microtubule-stabilizing docetaxel (DTX), a chemotherapeutic drug used to treat PCa, as well as other cancer types. HD-depleted cells showed significantly elevated plectin-dependent drug resistance at all concentrations studied (Figure 4I).

To gain insight into differentially regulated genes and cellular signaling pathways altered in PC3, α6-KO, β4-KO, PLEC-α6-dKO and PLEC-β4-dKO cells lines we performed an RNA-Seq analysis. In line with the functional analysis the most enriched pathways in PC3 α6-KO cells (PTEN/HD-negative) were related to proliferation and cell cycle regulation (Figures 4J-L). Curiously, except for the mitotic spindle pathway, all the most affected pathways were upregulated in plectin-dependent manner in β4-KO cells whereas only E2F targets and MTORC1 signaling were plectin-dependent in α6-KO cells indicating important differences between α6- and β4-KO cells (Figure 4L). To validate the transcriptomics data, we analyzed the expression of selected genes from the G2M_checkpoint pathway by qPCR. In agreement with the RNA-Seq data, *CCND1* and *CENPA* were upregulated in HD-deficient PC3 cells in plectin-independent manner (Figure 4M), while the increased expression of *MAD2L1* and *INCENP* was specifically abolished by PLEC-KO (Figure 4N). Taken together, our data suggest that plectin is a critical mediator of the tumorigenic properties of PTEN/HD-double negative PCa cells.

To assess the tumorigenic potential of selected cell lines *in vivo*, we analyzed the metastatic capacity of PC3, α6-KO and PLEC-α6-dKO cells using the SCID mouse tail-vein injection model. For this purpose, we first introduced eFFly luciferase and GFP expression into the different cell lines ^21^. Seven days after tail-vein injection into male ICR-SCID mice a luciferase signal indicated cell colonization of mainly the lungs (Figure 5A). Four weeks post injection, mice were euthanized and the average luminescence signal intensity in the lungs was measured. We observed that PC3 α6-KO cells colonized lungs much more efficiently than PC3 or PLEC-α6-dKO cells (Figures 5B-C). Immunofluorescence staining for human-specific phosphorylated keratin 8 confirmed the presence of human PC3 cells in mice lungs (Figure 5D). Interestingly, most of the tumors were localized in the upper part of the lungs, in close contact to cartilage, suggesting that PC3 cells, derived from a bone metastasis, prefer a harder surface for colonization (Figure 5C). To further confirm the metastatic capacity analysis, we also applied a novel bone marrow-on-chip assay. tdTomato-expressing MC3T3 osteoblasts and GFP-positive PC3 cells were co-cultured in collagen I-coated microfabricated microchannel chambers with a continuous flow of medium. Osteoblasts were first cultured alone for 7 days after which PC3 cells were added and allowed to colonize the chambers followed by a continued co-culture for 14 days (Videos S1, S2). PC3 cells were found to embed into osteoblast structures as relatively small foci while α6-KO cells formed much larger cell clusters (Figure 5F, Videos S3, S5). Surprisingly, addition of PC3 cells led to loss of osteoblasts in the co-culture while co-culture with PC3 α6-KO cells did not significantly affect osteoblast culture (Figure 5E). To analyze the drug-sensitivity of the different cells some of the co-cultures were treated with docetaxel starting at day 14 through day 21. PC3 cells displayed a clear dose-response to docetaxel treatment and were essentially eliminated after 7 days treatment with 1nM docetaxel (Figures 5E-G, Video S4). Of note, death of parental PC3 cells upon docetaxel treatment was accompanied by increase in the number of osteoblasts (Figure 5E). In contrast, PC3 α6-KO cells appeared highly resistant to docetaxel treatments as they seemed to be tightly integrated with the osteoblastic structures (Figures 5E-G, Video S6). This data suggests that PTEN/HD-negative cells (PC3 α6-KO) readily colonize osteoblast niches where they show robust resistance to docetaxel treatment when compared with parental HD-forming PC3.

**Figure 5.**
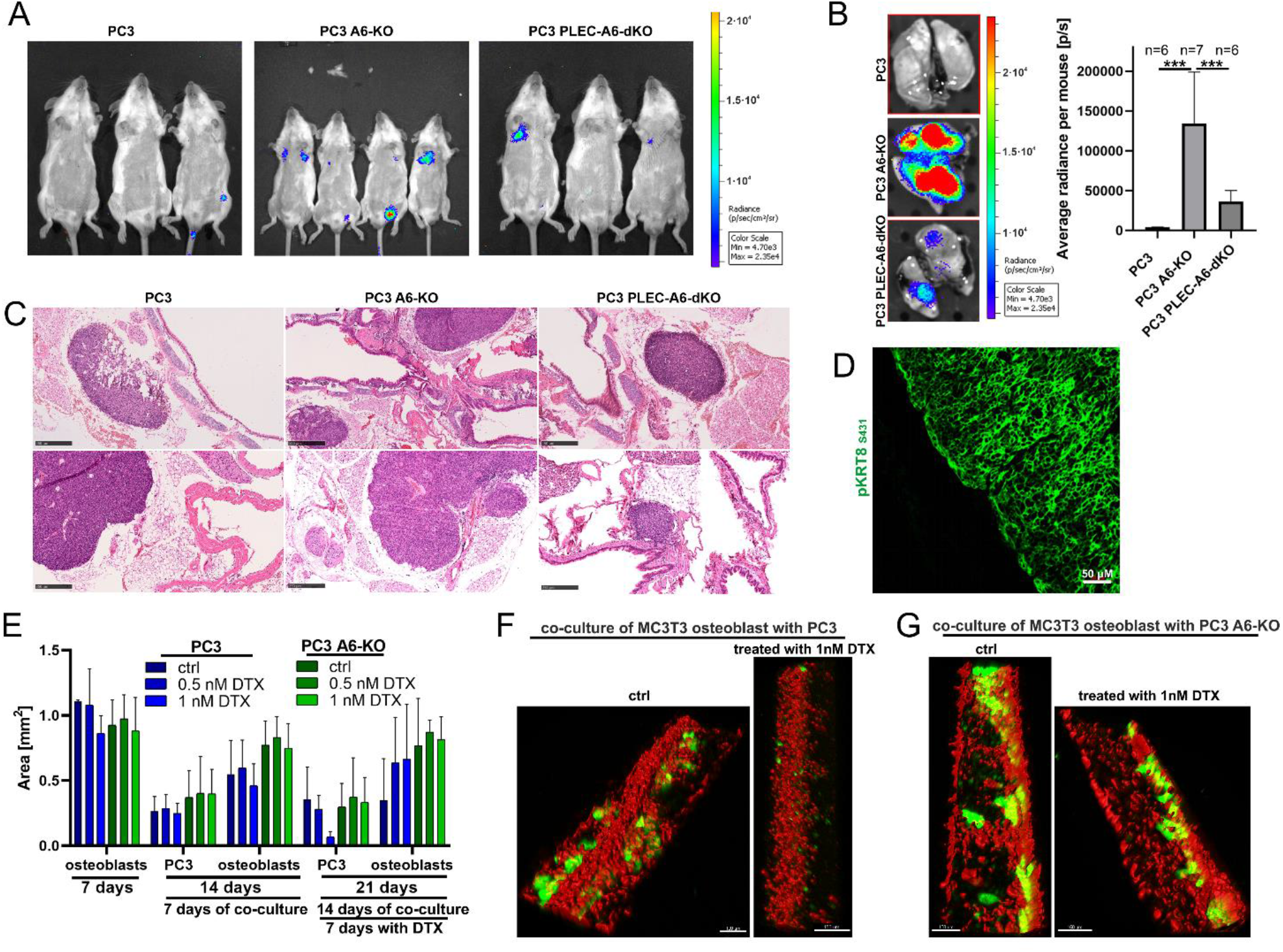
Loss of HDs promotes metastasis of PTEN-negative cells in vivo in a plectin-dependent manner. Luciferase activity signal in SCID mice 7 days after tail-vein injection of luciferase-GFP expressing PC3, PC3 α6-KO or PC3 PLEC-α6-dKO cells (A) a comparison of the luciferase activity signals in the lungs 4 weeks after injection (B); HE/EO staining of the lungs of SCID mice (C); immunofluorescence analysis of tumor foci in the lungs using an antibody recognizing human phosphokeratin-8 (D); bone marrow-on-chip analysis of the co-culture of Tdtomato-expressing MC3T3 osteoblasts and GFP-positive PC3 or PC3-α6KO cells treated or not with DTX (E); representative images of untreated and DTX-treated MC3T3 osteoblasts (red) / PC3 or PC3 α6-KO cancer cell (green) cocultures (F). For better visualization, osteoblasts were surface-rendered using IMARIS software.

### Dual HD- and PTEN-depletion is sufficient to transform non-invasive prostate cells and promote their metastatic capacity in vivo

Given the extensive genomic variability between different cancer cell lines we next wanted to confirm the causality of simultaneous loss of PTEN and HDs in promoting increased metastatic capacity by analyzing genetically engineered variants of the benign RWPE1 cell line for their metastatic capacity. For this purpose, RWPE1, RWPE1 β4-KO, RWPE1 PTEN-KO, RWPE1 α6/PTEN-dKO and RWPE1 β4/PTEN-dKO cells expressing the dual GFP-luciferase reporter construct were generated as above. To ensure detection of subtle differences between the metastatic capacity of the different variants only one hundred thousand cells of each population was injected into the tail-vein of male ICR-SCID mice. After four weeks, mice were sacrificed, blood and their lungs were collected. Histological analysis of lungs revealed micrometastases in the lungs of all the mice injected with RWPE1 PTEN-KO, RWPE1 PTEN-α6-dKO or RWPE1 PTEN-β4-dKO cells, and three out of six mice injected with RWPE1 β4-KO cells (Figure 6A). Very small microlesion was found in only one of the six RWPE1 cell injected mice. Importantly, cancer foci formed by PTEN/HD-dual KO cells were much bigger and more frequent in mouse lungs compared with foci formed with single PTEN-KO or β4-KO cells (Figure 6A). The human origin of the lesions in mouse lungs was confirmed by immunofluorescence staining using human phosho-KRT8-specific antibody (Figure 6B). The circulating cancer cells in blood were assessed using FACS sorting to separate the GFP-positive cancer cells from the blood. The amount of circulating PTEN-α6-dKO, PTEN-β4-dKO or β4-KO cells was significantly higher when compared with controls (Figure 6C). Higher levels of circulating cells depict increased survival capacity in the blood stream thereby potentially promoting their capacity to metastasize into distant organs. The harvested circulating cells were expanded and subjected to a western blotting analysis to confirm that isolated cells were still α6β4-deficient (Figure 6D). Remarkably, even the less abundant circulating parental RWPE1 and RWPE1 PTEN-KO cells recovered from the blood displayed strongly reduced levels of α6- and β4-integrins and relatively high levels of plectin levels compared to injected original cell populations (Figure 6D). Intrigued by this finding we assessed the functional properties of the cells recovered from blood. In line with our hypothesis, the ability of the recovered cells to migrate and invade strictly correlated with the expression levels of α6-and/or β4-integrins regardless of their original injected parental cell type (Figures 6E-F). When grown in 3D culture conditions, the recovered circulating cells formed cysts that generally resembled the phenotypes of the original injected cells although they appeared somewhat more unpolarized and disordered as was seen for example for α6-PTEN-dKO cysts that displayed elongated morphology with numerous protrusions (Figure 6G). In conclusion, our data indicates that dual HD-PTEN depletion transforms non-malignant cells, presumably via a plectin-dependent mechanism, enabling them to form micrometastases into lungs, as well as to survive in the blood stream.

**Figure 6.**
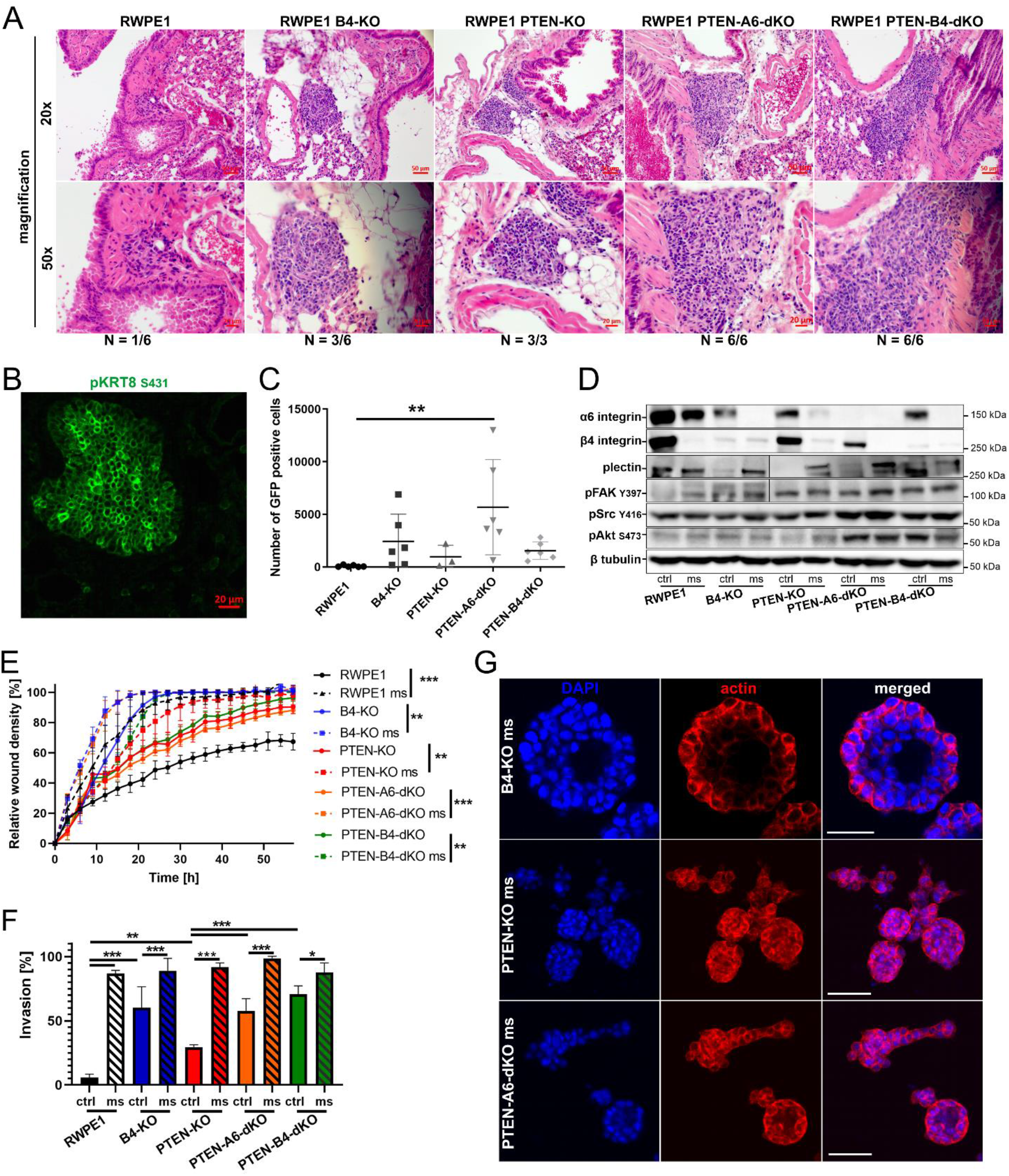
Dual loss of HDs and PTEN transforms nonmalignant RWPE1 cells into tumorigenic cells capable of in vivo metastasis into mouse lungs. HE/EO staining of lungs of the SCID mice with tail-vein injected RWPE1, RWPE1 β4-KO, RWPE1 PTEN-KO, RWPE1 PTEN-α6-dKO or RWPE1 PTEN-β4-dKO cells (A); immunofluorescence analysis of cancer foci with an antibody recognizing human phosphorylated keratin-8 (B); FACS analysis of the circulating GFP-positive RWPE1 cell variants in the blood of SCID mice (C); western blot analysis of the circulating RWPE1 cells recovered from the blood of SCID-mice (D); wound healing assay (E), invasion assay (F) and 3D culture (G) of recovered circulating RWPE1 cell variants. The data are presented as mean ± SD. All analyses were performed in triplicate.

### Upregulated plectin in PTEN- and HD-deficient PCa patients samples correlates with higher metastatic potential and worse patients’ survival

To investigate the clinical relevance of our findings we analyzed multiple patient cohorts available in public databases. Firstly, we checked the relative expression correlation between *PLEC* and *PTEN* levels in PCa. As shown in Figure 7A *PLEC* levels negatively correlates with *PTEN* levels. Secondly, based on the *PTEN* expression level we stratified the patients into *PTEN*-high and *PTEN*-low groups (Figures 7B-C). To further confirm the PTEN status we chose cohorts with information of *PTEN* copy number alterations and performed additional patient stratification based on *PTEN* copy loss into PTEN-normal and PTEN-loss groups (Figures 7J-R). We found that *PLEC* expression was significantly upregulated in both *PTEN*-low and *PTEN*-loss PCa groups (Figures 7B-C, 7J). We further stratified the *PTEN*-low and *PTEN*-loss groups according to their *ITGB4* or *ITGA6* expression levels and analyzed whether increased *PLEC* levels correlate with the different variables indicating PCa tumorigenesis. Higher *PLEC* expression levels were correlated with increased Prostate-Specific Antigen (PSA) levels, metastasis, Gleason score, tumor stage, and worse overall survival (Figures 7D-G, 7K-L). Importantly, high *PLEC* levels and lymph node metastasis, Gleason score, PSA level and metastasis all were positively correlated in double stratified *ITGB4* or *ITGA6* low, and *PTEN* low or *PTEN* copy loss number patients’ groups (Figures 7H-I, 7M-R).

**Figure 7.**
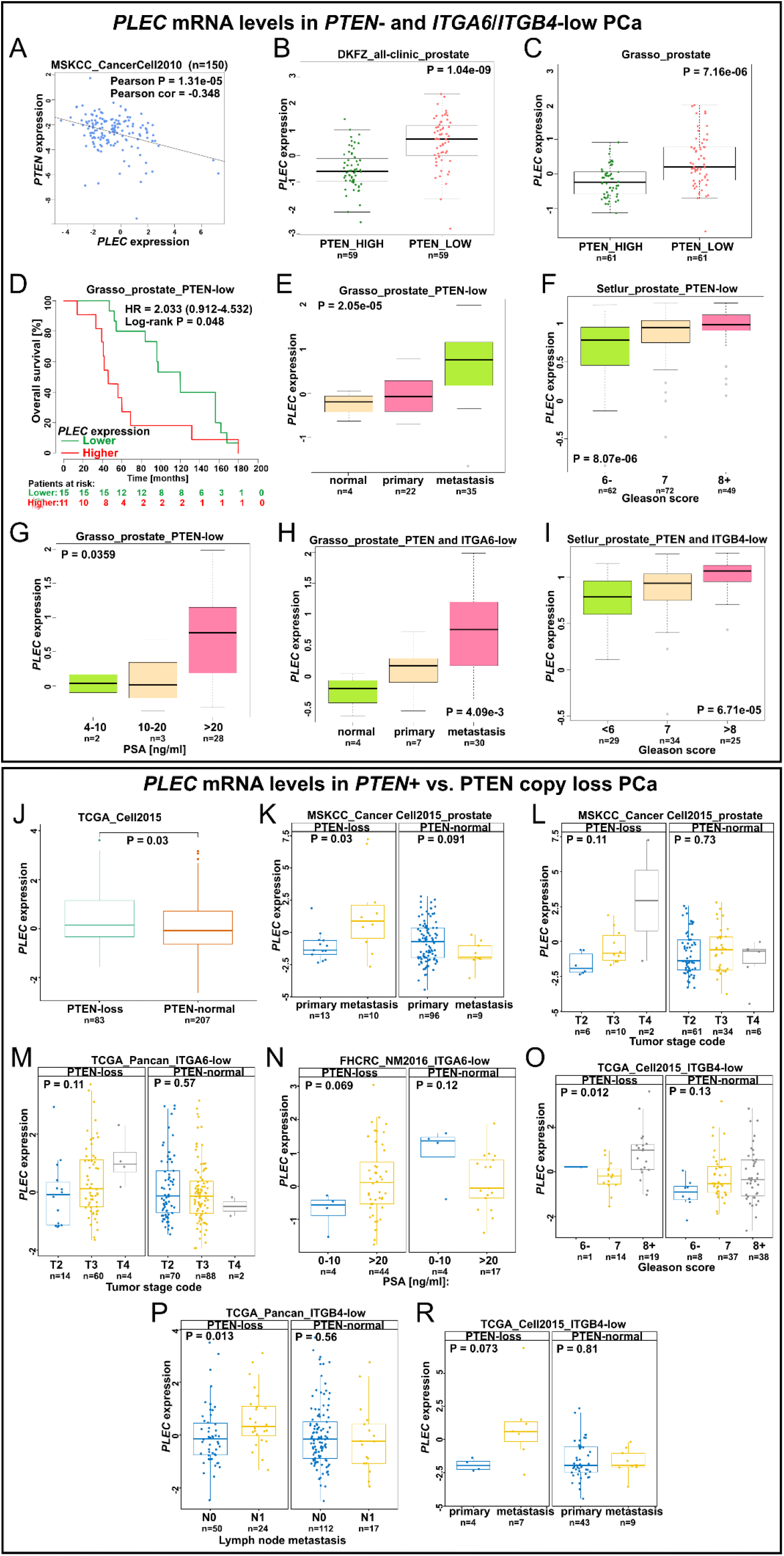
Upregulation of PLEC expression correlates with tumor aggressiveness in PCa patients with low PTEN expression and PTEN loss/deletion. Correlation analysis of PLEC-PTEN mRNA expression levels in PCa patients (A), comparison of PLEC level in PCa groups with high or low expression of PTEN (B-C); Higher PLEC level is associated with shorter overall survival (D) PLEC expression level is upregulated upon disease progression (E), higher Gleason score (F) and elevated PSA level (G) in PCa patients; PLEC level is upregulated in stratified PCa patients expressing low PTEN-ITGA6 (H) and PTEN-ITGB4 (I) level. Comparison of PLEC levels in PCa patients with PTEN loss/deletion and PTEN normal (wt) patient groups (J); PLEC is upregulated in metastasis samples (K) and advanced tumor stage (L) in PCa patients with PTEN loss. PLEC expression level is elevated in PCa patients with advanced tumor stage (M), elevated PSA level (N), higher Gleason score (O), lymph node metastasis (P) and metastasis (R) in stratified PCa patients with PTEN loss and expressing low ITGA6 or ITGB4 levels.

To validate bioinformatic data we generated a tissue microarray (TMA) from an independent PCa patient cohort consisting of 232 PCa tissue blocks. Four tissue cylinders were picked from each patient sample (two from normal or hyperplastic area and two from carcinoma regions). We found that HDs, as depicted by intense staining at the basal membrane of basal epithelial cells, were present only in normal glands and HD markers were either lost or presented more diffuse staining in carcinomas (Figure 8). Interestingly, in most PCa patients, β4 integrin expression was strongly reduced in tumor lesions whereas α6 integrin expression was retained although it displayed a diffuse staining and was evident in luminal cells. Plectin staining was similarly more diffuse in tumor lesions and the staining intensity was particularly high in carcinoma areas devoid of PTEN expression (Figure 8).

**Figure 8.**
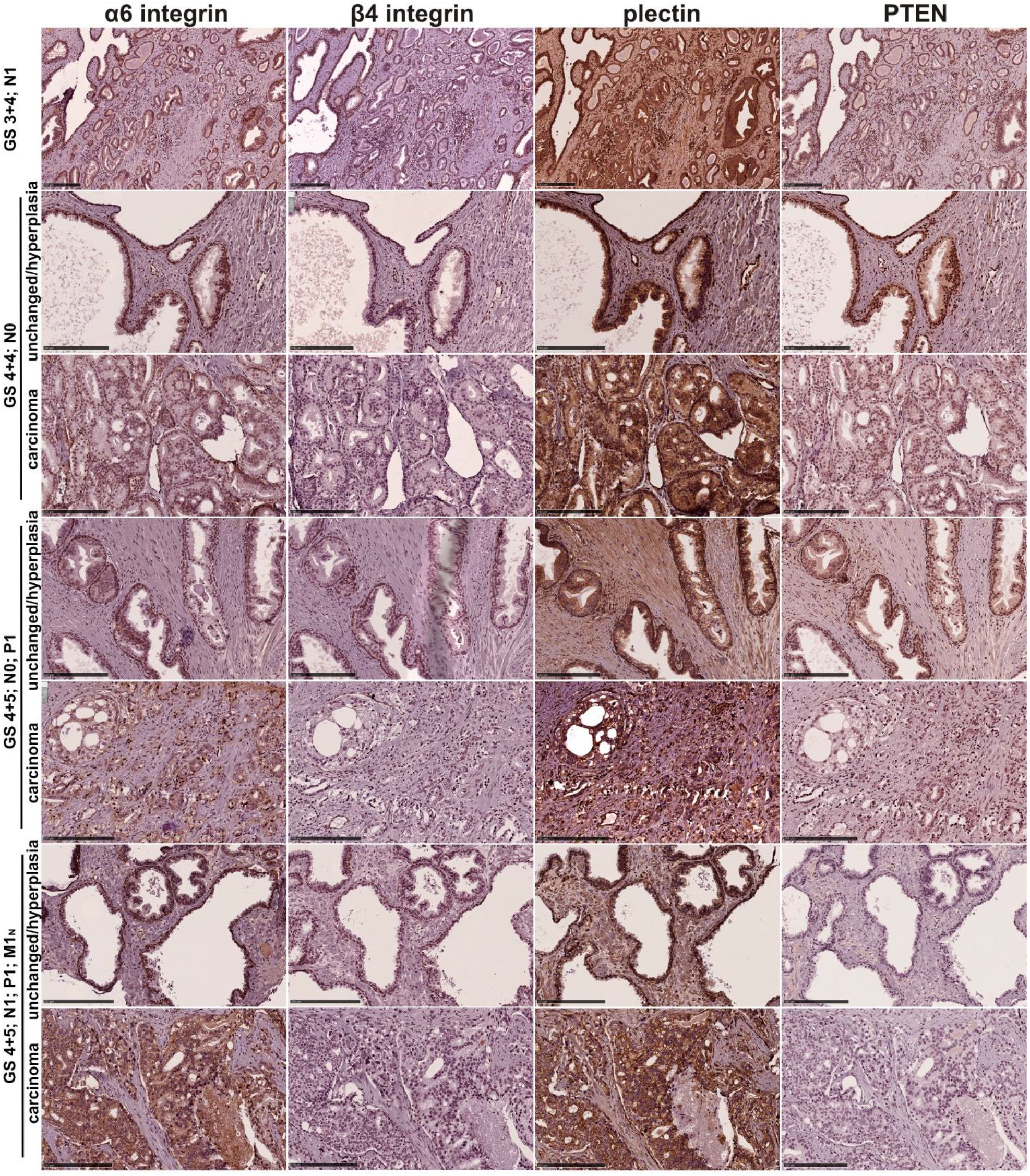
HDs are lost in prostate cancer and high plectin levels are observed in PCa cells with low levels of β4-integrin and PTEN. Representative images of α6-, β4-integrin, plectin and PTEN expression and localization in normal, hyperplastic and prostate carcinoma areas. GS – Gleason score, 1°+2°, N – node classification, P – progression, M – metastasis, M1_N_ – lymph node metastasis. In N,P,M: 0 – negative, 1 – positive. Scale bar 250 μM.

Collectively, analysis of several independent clinical datasets provides further evidence that plectin is a key tumorigenicity factor in PTEN/HD (α6 or β4)-double negative PCa tumors such that high *PLEC* expression co-occurring with low or absent *PTEN* and *ITGA6*/*ITGB4* expression correlates with higher PCa metastatic capacity, Gleason score, PSA level and worse overall survival.

## Discussion

Identification of critical molecular determinants driving the development of aggressive PCa remain to be identified. One of the frequently documented observations during PCa pathogenesis is the disruption of HDs, multiprotein assemblies formed around α6β4-integrins ^11–13^. HDs are dynamic structures regulating cell adhesion, differentiation, migration and invasion ^22^. Progressive loss of β4-integrin expression has been observed during the development of prostate intraepithelial neoplasia (PIN) into to invasive prostate carcinoma ^12^. However, the underlying mechanisms how HDs modulate tumorigenesis remain elusive. Here, we report that HDs are reorganized or lost in a broad panel of prostate cell lines further corroborating disruption of HDs as one of the most critical events during PCa progression. In functional studies, we established a causal link between loss of HDs and tumorigenesis in PTEN-negative PCa cells. This new finding is highly relevant because abrogation of the tumor suppressor function of PTEN is one of the most frequent genomic aberrations in PCa patients ^18^. In line with our data, recent study by Dalton et al. showed that transcriptional corepressor C-terminal binding protein 1 (CTBP1), which is overexpressed in prostate cancer, downregulates several genes relevant to cell adhesion, including *ITGB4*, in PTEN-negative PC3 cells ^23^. It was shown that reduced *ITGB4* expression in PC3 cells was critical for efficient metastasis in NOD-Scid Gamma mice. Using an extensive CRISPR/Cas9-mediated genetic engineering approaches in different cell lines, we demonstrated that simultaneous loss of PTEN and HDs (α6-KO or β4-KO) in prostate epithelial cells induced several tumorigenic properties including proliferation, migration, anoikis-apoptosis- and drug-resistance *in vitro*, as well as increased metastatic capacity *in vivo*. Mechanistically, dual PTEN/HD depletion strongly activated the EGF/PI3K/Akt- and FAK/Src-pathways. Importantly, all the above-mentioned pro-tumorigenic effects were to large extent abolished by plectin downregulation indicating a key role for plectin in these processes.

Plectin is a key component of HDs as it binds to the cytoplasmic tail of β4-integrin and links it to the cellular IF network. Plectin is a large (∼500 kDa) versatile cytoskeletal linker molecule capable of binding also to the actin and microtubule networks. It is abundantly expressed and can exist as several alternatively spliced isoforms. Upregulated plectin levels have been correlated with the progression of prostate as well as other cancers ^24–27^. Moreover, a recent study implicated plectin as the key regulator of PCa growth and metastasis ^28^. Our data is in line with these findings and reveals that loss of PTEN expression was required to stabilize or upregulate plectin levels upon HD disruption. In contrast to PTEN-deficient cells, disruption of HDs (α6-KO or β4-KO) in normal epithelial cells led to robust downregulation plectin. Importantly, this finding was not only restricted to PCa cells as it was also seen in breast cancer cells. Thus, our data suggest that plectin is specifically a critical mediator of the tumorigenic potential of PTEN/HD-double negative PCa cells.

Curiously, in α6- or β4-integrin depleted prostate epithelial cells, plectin displayed partial colocalization with FAs linking to the actin cytoskeleton, especially in PTEN-negative cell lines. This was accompanied by stimulated integrin signaling and cell migration. Competitive binding of α6β4-integrin and actin to plectin has been reported ^29^. More recently, HD-deficient keratinocytes were reported to show enhanced FA maturation and upregulated FAK signaling ^30^. Together, these observations propose a model where plectin preferentially associates with the IF-linked HDs but upon HD disruption switches to actin-linked FAs facilitating signaling cascades promoting cell migration and growth. Further studies are warranted to investigate such potential switch in more detail, for example to assess whether different plectin isoforms are involved.

Interestingly, loss of α6- or β4-integrin expression in normal prostate epithelial cells (RWPE1) caused a strong reduction of their heterodimer partner (and of plectin), whereas in PTEN-negative cells the heterodimer partner expression was only modestly affected and plectin expression was maintained, or even upregulated, suggesting that the retained subunit might play an additional functional role in HD-depleted cells. Given that the plectin binding site in the α6β4-heterodimer locates to the cytoplasmic tail of β4-integrin, it would be interesting to study differences of the cellular signaling outcome depending on whether α6- or β4-subunit expression is retained. In the absence of β4-integrins, α6-integrin may pair with β1-integrins whereas no alternative α-partner has been documented for β4-integrin, although there is evidence that β4-integrins can interact with several HD components and reach the cell surface also in the absence of α6-subunit ^31, 32^. It is noteworthy, that while the overall phenotype of α6-KO and β4-KO cells in the current experimental settings were similar, there were also some context-dependent differences such as cell migration in the PTEN^+^ background (Figure 3D) or transcriptional effects on the proliferation and growth-associated pathways (Figure 4). It is noteworthy, that while the loss of HDs was found to be causal, the loss of β4-(229 out of 232) rather than α6-integrin was observed in most PCa patients even in PTEN-positive background. In line with this observation, bioinformatic analysis revealed that *ITGB4* reduction correlated with higher PCa metastatic capacity in PTEN independent manner while *ITGA6* levels were significantly correlated only in PTEN copy loss number group. More detailed studies are needed to address the specific functional roles of α6- and β4-subunits upon HD disruption.

PTEN-HD dual negative PCa cells showed increased numbers of circulating tumor cells (CTC) in the *in vivo* metastasis model. The role of CTC is still somewhat controversial, but promising reports on the potential of CTCs as refined prognostic markers helping to identify personalized treatments for patients suffering from various cancer types have been published ^33–35^. CTCs have been proposed to predict the survival rate, diagnosis and stage of pancreatic adenocarcinoma (PDAC) ^36, 37^. Recently, plectin was proposed to be a biomarker for CTCs of PDAC improving differentiation between malignant pancreatic disease and chronic pancreatitis ^38^. Here, we observed that PTEN-HD dual negative cells displayed a high number of CTCs and robust plectin expression. Moreover, CTCs in general appeared to have relatively higher levels of plectin expression when compared with their injected parental cells, inversely correlating with α6-and/or β4-integrin expression. Our data is thus in agreement with a potential role of plectin as a valuable CTC-marker predicting the metastatic potential of CTCs.

Taken together, we identify and characterize a novel mechanism where dual loss of PTEN expression and HD assembly synergistically drive PCa progression by inducing plectin-dependent activation of Src/FAK- and EGFR/PI3K-signaling pathways which in turn promote cell migration and growth *in vitro* and metastatic capacity *in vivo*. These data were corroborated by extensive bioinformatic analysis of multiple clinical PCa data sets as well as independent PCa patient validation cohort analyses. Due to strong correlation between PTEN-HD loss and PCa severity, our data not only reveals new potential therapeutic targets, but also suggests novel diagnostic analyses that may help to stratify PCa patients by identifying those individuals with high risk for development of aggressive disease.

## Materials and methods

### Cell culture

RWPE1, DU145, PC3, 22Rv1, LNCaP, VCaP and JIMT-1 cell lines were originally purchased from ATCC. LNCaP 1F5 and V16A were a gift from Dr. Olli Jänne from the University of Helsinki. RWPE1 were cultured in Keratinocyte SFM medium (Gibco) supplemented with bovine pituitary extract and human recombinant EGF, and standard antibiotics: penicillin (100 units/mL) and streptomycin (100 μg/mL), according to manufacturer’s protocol. PC3 and DU145 cells were maintained in F12K (Gibco) and MEM (Gibco) medium, respectively containing 10% fetal bovine serum (Gibco) and standard antibiotics. 22Rv1, VCap, LNCap, LNCap IF5, V16A and JIMT-1 cells were grown in RPMI1640 (Sigma-Aldrich) containing 10% FBS and standard antibiotics. The cells were maintained at 37°C in a humidified atmosphere with 5% CO2. All cell lines were confirmed to be mycoplasma-free during the analysis.

### Plasmid construction

To generate knock-out cells the specific gRNA sequence was inserted via *BsmB*I into plentiCRISPRv2 vector as follows: for α6-integrin knock out: TTTTCTTTGGACTCAGGGAAAGG (exon 6), for β4-integrin: CTGCACGGAGTGTGTCCGTG (exon 3) and CAACTCCATGTCCGATGATC (exon 5), for plectin: TGAGGTTGTGGCCATCGCGG (exon 3), TCATACAGCGACGAGACGT (exon 8) and GGACGCATCCGCAGCAACG (exon 31) were applied. For α6-integrin overexpression, the CDS sequence was amplified from cDNA extracted from PC3 cell line and inserted via *Sal*I into pBABEhygro vector. To recover α6-integrin expression in knock-out cells (α6-KO), using site-directed mutagenesis the single nucleotide substitution in PAM sequence was inserted in pBABEhygro_*ITGA6* (Gly=>Ala). All of the plasmids were verified by sequencing.

### Viral transduction

Lentiviral particles were produced by co-transfecting 5 μg of the plentiCRISPRv2 containing specific gRNA sequence and 3.75 μg psPax2, and 1.25 μg pVSV-G (second-generation lentivirus packaging system) into human embryonic kidney 293T cells (ATCC, LGC Standards GmbH, Wesel, Germany; CRL-11268) using Lipofectamine 2000 (Invitrogen, Thermo Fisher Scientific). To generate retroviral supernatant 4 μg pBabe-hygro based plasmid and 0.5 μg pVSV-G were co-transfected into Phoenix gag-pol cells (www.stanford.edu/group/nolan/retroviral_systems/phx.html) ^39^, obtained from ATCC with authorization by Garry Nolan, School of Medicine, Stanford University, Stanford, CA). The transduced cells were selected with 750 μg/mL of G418 (geneticin) (Gibco, Paisley, UK) or 1 μg/mL puromycin or 50 μg/mL hygromycin B for at least 7 days and subsequently analyzed by western blotting.

### Western blotting

Cells were grown to reach 80–90% confluency and then washed in PBS (Gibco) and scraped in RIPA buffer: 10 mM Tris-HCl pH 8.0, 150 mM NaCl, 0.5% SDS, 1% IGEPAL, 1% sodium deoxycholate containing 2 mM PMSF (phenylmethylsulfonyl fluoride), 10 μg/mL aprotinin, and 10 μg/mL leupeptin. Protein concentration was estimated using BCA Protein Assay Kit (Pierce). 30 ug of protein lysate was resolved by SDS-PAGE and transferred onto a Protran pure 0.2 micron nitrocellulose (Perkin Elmer). The membranes were incubated for 1h in 5% skimmed milk and probed with specific primary antibodies (Table S1) overnight at 4 °C. Secondary antibodies conjugated with HRP and Lumi-Light Western Blotting Substrate (Roche) were used to visualize specific protein bands. The bands were detected using Fujifilm LAS-3000 bioimaging and scientific research imaging equipment (FUJI PHOTO FILM CO.,LTD.).

### Quantitative RT-PCR

RNA was extracted from the cells using RNeasy Mini Kit (QIAGEN) according to the manufacturer’s protocol. The RevertAid reverse transcriptase (Thermo Scientific) was used to synthesize cDNA from 1 ug RNA and Brilliant III Ultra-Fast SYBR Green QPCR Master Mix (Agilent Technologies) was used for the Quantitative RT-PCR reactions. At least three replicates in at least two independent experiments were applied for each gene and the data were normalized against GAPDH (control). Primer sequences used in this study are collected in Table S2.

### Proliferation assay

2·10^3^ of PC3, DU145, 22Rv1 and JIMT-1 or 3·10^3^ of RWPE1 cells were seeded onto 96-wells plate in a culture medium. The area of proliferating cells in time was analyzed using IncuCyte S3 Live-Cell Analysis System (Essen Bioscience Inc.).

### Wound healing assay

6·10^4^ cells were seeded on Incucyte ImageLock 96-well Plate (Essen BioScience Inc. 4379) and cultured 24-48h to reach full confluency. The wound was done using the Woundmaker 96 tool (Essen Bioscience Inc.) and the migration of the cells to heal the scratch was analyzed using IncuCyte S3 Live-Cell Analysis System (Essen Bioscience Inc.).

### Invasion assay

100 μl of diluted in prechilled serum-free medium Matrigel matrix basement membrane (0.2 mg/mL) (Corning 354230) was loaded into 8-μm TC inserts (Greiner Bio-One) and allowed to solidify in 37°C for 1h. In the next step, 5·10^4^ RWPE1 or 2.5·10^4^ PC3 cells in maximum 180 μl total volume were gently seeded on the top of the gel. The lower chambers were filled with 600 μl of medium containing 10% FBS as an attractant. After 24 or 48h for PC3 and RWPE1, respectively, the cells were fixed using 4% PFA in PBS and stained with 0.02% crystal violet solution in 10% ethanol. Noninvasive cells in the matrix layer were discarded using a cotton swab and the cells on the external surface of the inserts were analyzed in the bright field using Zeiss Axio Vert.A1 microscope.

### 3D culture

150 μl of Matrigel matrix basement membrane (Corning 354230) was evenly spread on 35 mm glass-bottom μ-Dish (IBIDI) and allowed to solidify at 37°C for 1h. 5·10^3^ cells were seeded on the top of Matrigel matrix overlay and cultured in standard medium containing 2 % Matrigel matrix for 7 days. For RWPE1 medium was additionally supplemented with 5 % FBS. After this time, the cells were gently washed twice with PBS, fixed with 4% PFA with 0.1 % glutaraldehyde and followed standard immunofluorescence protocol. The organoids were analyzed by using Olympus FluoView FV1000 confocal microscope.

### Immunofluorescence microscopy

The cells growing on 35 mm glass-bottom μ-Dish (IBIDI) were gently washed twice with PBS and fixed with 4 % PFA in PBS for 15 minutes. To quench unspecific PFA fluorescence, cells were incubated with 100 mM glycine in PBS for 20 minutes and then permeabilized with 0.1 % Triton X100 in PBS for 15 minutes. After this time, the samples were blocked with 0.2 % gelatin, 0.5 % BSA in PBS for 1 h and then incubated with primary antibody diluted in blocking buffer overnight at 4°C (Table S1). In the next step, the samples were washed 4 times with blocking buffer and probed with secondary antibody conjugated with a fluorophore for 1 h or overnight in 4°C. After the same washing with PBS, the cells were analyzed using Zeiss LSM 780 confocal microscope or Zeiss Cell Observer.Z1 Spinning Disc confocal microscope. ZEN Blue software was applied to data analysis.

### Soft agar assay

The 6 well plate was coated with 1.5 ml 0.7% SeaPlaque agarose solution dissolved in a standard culture medium. After solidification, 5·10^3^ cells were suspended in 1 ml of 0.4 % SeaPlaque agarose in culture medium and seeded on the lower layer and covered with 200 ul culture medium. Cells were cultured for 21 days at 37°C and 5% CO_2_, and culture medium was exchanged twice per week. The colonies were stained with 0.01% crystal violet in 10% ethanol and captured by Leica MZ6 stereo microscope. ImageJ software was applied to analyze the size of colonies and the total area of colonies.

### PolyHEMA assay

The 6-wells plate was coated with 70 μl of 20 mg/mL polyHEMA solution in 96% ethanol overnight in RT. In the next step, 4 ·10^4^ cells in standard medium were seeded on the plate. At selected time points the floating cells were collected, washed with cold PBS and stained with Annexin V-fluorescein isothiocyanate/propidium iodide kit (BD Pharmingen). The number of apoptotic cells was determined using BD Accuri Flow Cytometer.

### MTT assay

4·10^3^ cells were seeded on 96-wells plate and allowed to grow overnight. Next day the medium was replaced with a fresh one, containing selected docetaxel concentration and cells were cultured for the next 72h. Since the drug was dissolved in DMSO, the maximum volume of DMSO corresponding to maximum docetaxel dose was applied as a control. The survival was assessed by incubation with 50 ul of 4mg/ml MTT (3-(4,5-dimethylthiazol-2-yl)-2,5-diphenyltetrazolium bromide) in PBS for 4 h. The formazan crystals were dissolved in 100% DMSO, and absorbance was measured at 570 nm using VICTOR3 Multilabel Plate Reader (Perkin Elmer).

### RNA seq

6 ·10^4^ cells were seeded for 36h on 6-wells plate coated with 10 μg PureCol Type I bovine collagen (Advanced BioMatrix). The total RNA was extracted using the RNeasy Mini Kit (QIAGEN) according to the manufacturer’s protocol. The concentration and quality of RNA samples were determined using a NanoDrop 2000 micro-volume spectrophotometer (Thermo Scientific). The library construction and sequencing were performed by Novogene Europe (United Kingdom). Paired-end, 150 bp read-length sequencing was carried out using Illumina PE150 Novaseq and 6 Gb reads were generated for each sample. Raw sequence reads were first pre-processed with FastQC ^40^ for quality control. Trimmomatic ^41^ was employed to process reads for quality trimming and adapter removal. A final FastQC run on cleaned reads was conducted to ensure the read quality. The processed reads were aligned against the human genome assembly hg38 using STAR version 2.7.2a ^42^ with default settings. HTSeq (htseq-count) was employed to quantitate aligned sequencing reads against gene annotation from Encode and with parameters “-s no, –i gene_name”. Differential expression analysis was performed from read count matrix using Bioconductor package DESeq2 (1.26.0) ^43^. Genes with low expressions (< 2 cumulative read count across samples) were filtered out prior to differential expression analysis. A threshold of FDR < 0.1 was applied to generate the differentially expressed gene list. Data was normalized using variance Stabilizing Transformation (VST) method from DESeq2. Heatmap displaying gene expression levels was generated using R package “pheatmap” (1.0.12).

### Gene Set Enrichment Analysis

Functional annotation of gene expression profiles was performed using Gene Set Enrichment Analysis (GSEA). The “stat” statistics from the differential gene list was sorted in a descending order for the generation of the pre-ranked gene list. GSEAPreranked test ^44^ was applied to examine the pathway enrichment from Hallmark in MSigDB database. All parameters were kept as default except for Enrichment statistic = “weighted”, Max size (exclude larger sets) = 5000, number of permutations =1000.

### In vivo mouse model

6-7 weeks old IcrTac:ICR-Prkdc (SCID) mice (male) (Taconic Biosciences A/S) were injected with 1·10^5^ of RWPE1 or 1·10^6^ of PC3 cells in 250 μl (total volume) of PBS into the tail vein. For every group, except RWPE1 PTEN-KO (3 mice died during the procedure) 6 mice were applied. After 4 weeks, mice were sacrificed. 0.5-0.6 ml of blood was immediately collected from the aorta. Lungs were analyzed macroscopically, fixed in 4 % PFA in PBS for 48h and then washed in water and transferred to 70% ethanol for 48h, embedded in paraffin and followed hematoxylin and eosin staining. Immunocompromised mouse lines were maintained under internal permissions from the Laboratory Animal Centre of University of Oulu (numbers 33/2021, 34/2021). Mouse cancer models and protocols are approved by the National Animal Experiment Board (ESAVI/3901/2021). In all animal work, the principle of 3R (reduction, refinement, replacement) is respected.

### Circulating cells recovery

The blood collected from mice was immediately diluted in RBC lysis buffer (0.8% NH_4_Cl, 0.084% NaHCO_3_, 0.037% EDTA), mixed for 5 minutes, washed in PBS and the cells were seeded on 10 cm plate in culture medium. After 2 days the cells were harvested by trypsinization, washed in PBS and analyzed using BD FACS Aria Illu (BD Biosciences). GFP-positive cells were recovered from the cell mixture, seeded on a new plate and cultured in the standard medium.

### Immunohistochemical staining

The tissue slides were deparaffinized by incubation for 1h at 55°C. To rehydrate, twice washing in xylene followed 100% ethanol, 94% ethanol and 70% ethanol (5 minutes in each) was applied. Heat mediated antigen retrieval was performed in the next step, by incubating the slides in boiling citric acid buffer (pH 6.0) for 10 minutes, then washed in PBS, incubated with 3% H_2_O_2_ for 10 minutes and then blocked with 5% BSA in PBS for 1h. After overnight incubation with primary antibodies at 4°C, the slides were washed in PBS and probed with HRP-conjugated secondary antibody for 1h. The positive antibody signal was revealed using DAB Substrate Kit (Abcam, ab64238). After staining in Harris hematoxylin solution (2 minutes) and dehydration with 70% ethanol, 94% ethanol, 100% ethanol and twice xylene (5 minutes in each) the slides were analyzed using Zeiss Axio Imager.M2 microscope.

### Bone marrow-on chip assay

Bone marrow-on-chip was fabricated in polydimethylsiloxane (PDMS) (Momentive, UK) using replica molding. The chip was molded from 10:1 ratio (by weight) of base to curing agent, and cured for 4 hours at 65°C. The two parts are then bonded to a glass slide with PDMS slurry and allowed to bake at 65°C for 4 hours. The fabricated chip contains 24 separate units (each 50 µL volume), in which 4 parallel microchannels are connected with 2 reservoirs. The channel widths tapers from 1000 µm to 100 µm between the opposing inlets. The chips were rinsed with 96% ethanol and UV sterilized before use. Next, the microchannels were coated with collagen I for 1h at 37°C on a lab rocker (1 cycle/15 minutes). 2.5·10^4^ stably Tomato expressing-MC3T3 osteoblasts were loaded into the inlet of each microchannel, filled with α-MEM medium (Lonza) supplemented with 10% FBS, 8 mg/ml ascorbic acid and standard antibiotics and cultured at 37°C, 5% CO_2_ on a lab rocker to provide a proper fluid flow (1 cycle/15 minutes). After 7 days, osteoblasts were imaged using Z-stack acquisition in Leica SP8 Falcon confocal microscope in live-cell chamber maintaining 37°C and 5% CO_2_. and then, 2.5·10^4^ stably GFP-expressing PC3 cells were added to the osteoblasts and co-cultured in α-MEM medium (Lonza) supplemented with 10% FBS, 8 mg/ml ascorbic acid and standard antibiotics for next 7 days in the same conditions. Co-culture of PC3 cells and osteoblasts was assessed using Leica SP8 Falcon confocal microscope in a live-cell chamber maintaining 37°C and 5% CO_2_ and after this 0.5 nM or 1 nM DTX was added to the medium. As a control used DMSO corresponding to maximum docetaxel volume. After next 7 days, the same positions on the plate were imaged to analyze the effect of DTX on cell survival. The data were analyzed using IMARIS x64 9.2.1 software. The comparison of the area of MC3T3 cells calculated using IMARIS software is presented in Video S1 and S2.

### Tissue microarray

TMA is derived from a cohort of 232 men operated during 2005-2017 in Hospital of Oulu, and living in Oulu region to have the follow-up data available. Most suitable formalin-fixed paraffin-embedded (FFPE) tissue block was selected from 232 cases. Annotations for TMA construction were marked in hematoxylin and eosin-stained slides or digitized slides. Two cores from tumor tissue and two cores from benign tissue were obtained using 2 mm punches. The TMAs were constructed with Galileo TMA CK4500 microarrayer (Isenet, Milan, Italy) in collaboration with the biobank Borealis (Oulu University hospital). The use of pathological archive material was licensed by the National Supervisory Authority for Welfare and Health. This study was approved by the Ethics Council of the Northern Ostrobothnia Hospital District.

### Bioinformatic analysis

RStudio (v. 1.2.5033) with R (v. 3.6.3) was used for statistical analyses ^45, 46^. Data were retrieved from the cBioPortal for Cancer Genomics database and Oncomine ^47–49^. Mann-Whitney U test and Kruskal-Wallis H test were used to assess statistical significance between two or more groups, respectively. The expression correlation analysis was assessed by Pearson’s product-moment correlation. While PCa patients without PTEN copy number alterations were grouped as PTEN normal, PTEN loss was defined as patients with PTEN shallow or deep loss. In another stratification method, patients were stratified to high or low groups based on the median gene expression levels. Kaplan Meier survival analysis was performed to investigate the association between gene expression levels and patient prognosis by using R package “Survival” (v. 3.2.3) and “Survminer” (v. 0.4.7) ^50–52^. Statistical analyses for all Kaplan Meier curves and hazard ratios (HR) were calculated using log-rank test and cox proportional hazards model, respectively ^53^. Samples with missing data were excluded from analyses. *P* value < 0.05 was considered statistically significant.

### Statistical analysis

Data are expressed as means ± SD of at least three independent experiments. Comparative data were analyzed with the unpaired or paired Student’s *t*-test or Two-way ANOVA using GraphPad Prism 9 software. The results were considered statistically significant when the *p*-value was less than 0.05 (*), 0.01 (**), or 0.001 (***).

## Acknowledgments

We thank Riitta Jokela for overall expert technical assistance, Jaana Träskelin for expert technical assistance at Biocenter Oulu Virus Core Laboratory, Dr Veli-Pekka Ronkainen for expert assistance in microscopy at Biocenter Oulu Tissue Imaging Center and Dr Virpi Glumoff for expert assistance in FACS. This work was funded by University of Oulu and Academy of Finland PROFI3 program and Jane and Aatos Erkko Foundation and Biocenter Finland.

## Author contributions

Conceptualization: AM; Methodology: TW, RD; Validation: TW, AS, XY; Formal Analysis: TW, QZ; Investigation: TW, AS; Resources: AA, MV, QZ, RD, PS; Writing – Original Draft: AM, TW; Writing – Review & Editing: AM, TW, G-HW, QZ, RD, AA, MV; Visualization: TW, AS, QZ, AM; Supervision: AM; Funding Acquisition: AM

