## Supplemental data for "Disassembly of hemidesmosomes promotes tumorigenesis in PTEN-negative prostate cancer by targeting plectin into focal adhesions"

### Extended Supplemental Data for Wenta et al. Disassembly of hemidesmosomes promotes tumorigenesis in PTEN-negative prostate cancer by targeting plectin into focal adhesions

### Supplemental figures

*Figure S1. Immunofluorescence analysis of β4-integrin in RWPE1 α6-KO and PC3 α6-KO cells.*

*Figure S2. Rescue of α6-integrin expression in α6-deficient (α6KO) cells restores their enhanced migratory ability down to normal level.*

*Figure S3. Dual depletion of HDs and PTEN increases proliferation and migration of JIMT-1 - breast cancer cells.*

*Figure. S4 Downregulation of ITGB4 or ITGA6 correlates with tumor aggressiveness in PCa patients with low PTEN expression.*

*Figure S5. Simultaneous loss of PTEN and ITGB4/ITGA6 reduction increases tumor aggressiveness in PCa patients.*

**Supplemental tables**

*Table S1 List of antibodies used in this study.*

*Table S2 List of qPCR primers.*

**Supplemental videos**

*Video S1. Area of MC3T3 osteoblasts grown for 7 days - imaged using Leica SP8 Falcon microscopy.*

*Video S2. Surface-rendered presentation of video S1 (MC3T3 osteoblasts) using IMARIS software.*

*Video S3. Co-culture of MC3T3 osteoblast with PC3 cancer cells – control (21d).*

*Video S4. Co-culture of MC3T3 osteoblast with PC3 cancer cells (21d) treated for 7 days with 1nM DTX.*

*Video S5. Co-culture of MC3T3 osteoblast with PC3 α6-KO cancer cells – control (21d).*

*Video S6. Co-culture of MC3T3 osteoblast with PC3 α6-KO cancer cells (21d) treated for 7 days with 1nM DTX.*

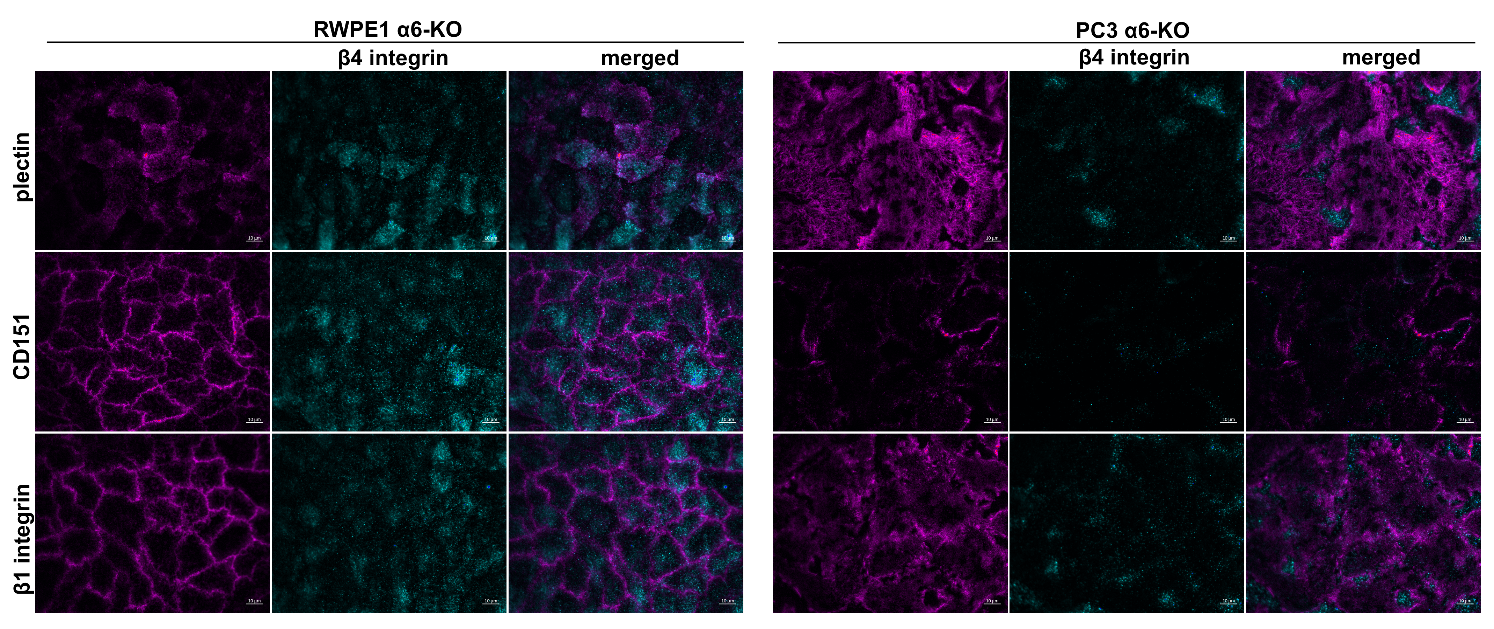
***Figure S1. Immunofluorescence analysis of β4-integrin in RWPE1 α6-KO and PC3 α6-KO cells****. Localization of β4-integrin (cyan) in the absence of α6-integrin expression was determined by immunofluorescence analysis in relation to plectin (magenta, top row), CD151 (middle row) and β1-integrin (bottom row).*

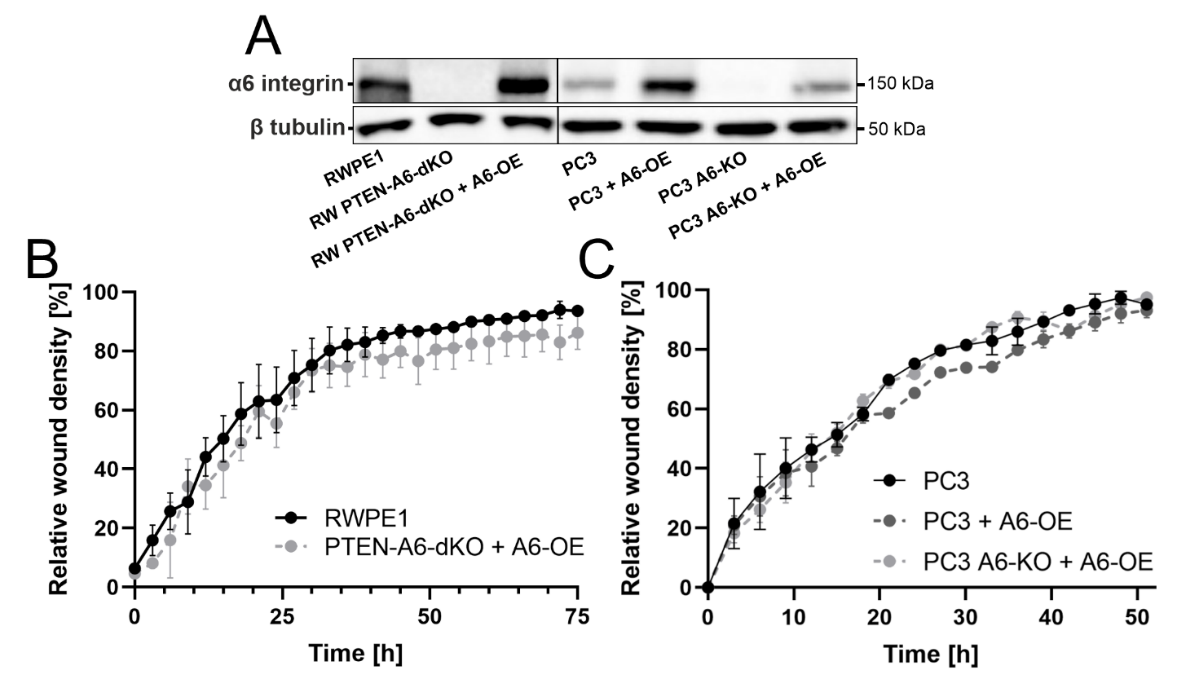

***Figure S2. Rescue of α6-integrin expression in α6-deficient (α6KO) cells restores their enhanced migratory ability down to normal level****. Western blotting analysis was done to confirm rescue of α6-integrin expression (A); wound healing assay for RWPE1 PTEN-α6dKO with recovered α6 integrin expression (B) wound-healing assay for PC3 α6-KO with recovered α6 integrin expression and PC3 with α6 integrin overexpression (C).* *The data are presented as mean ± SD. All analyses were performed in triplicate.*

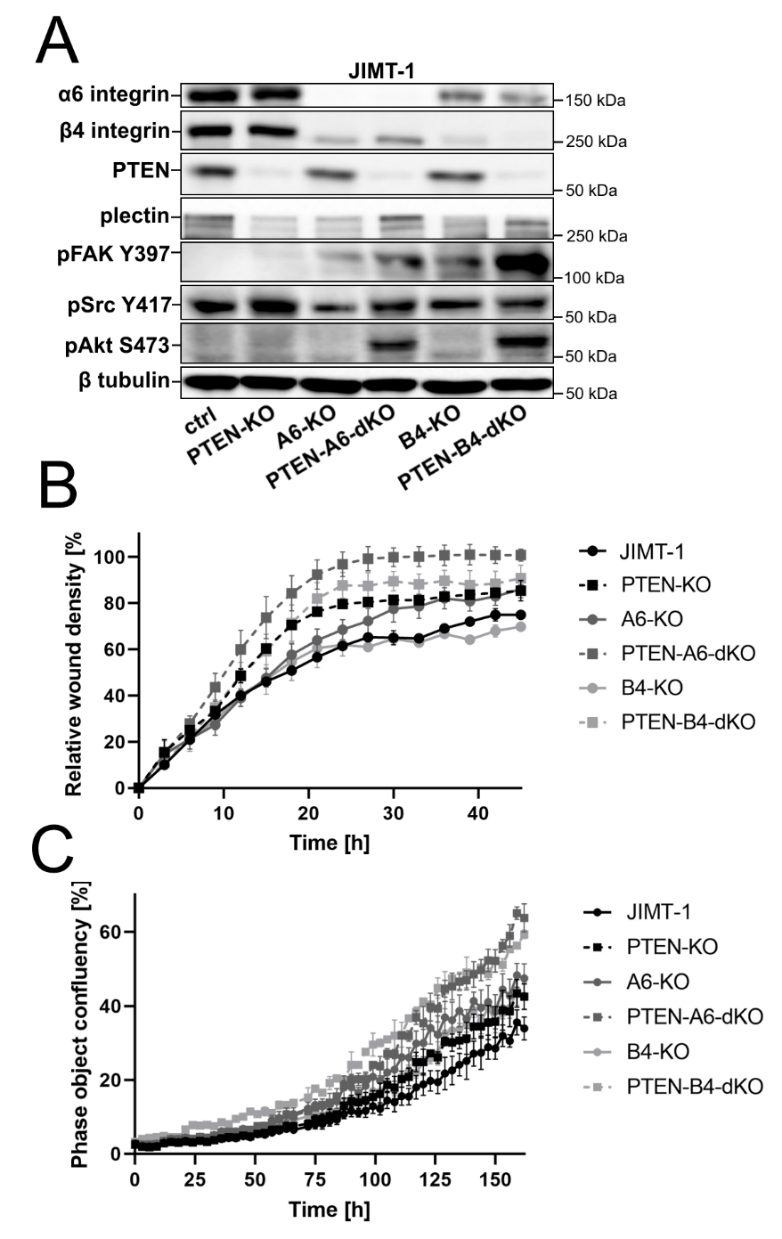

***Figure S3. Dual depletion of HDs and PTEN increases proliferation and migration of JIMT-1 - breast cancer cells****. Western blotting analysis of JIMT-1 cell variants with HD- and PTEN-depletion (A); Wound healing assay (B) and proliferation assay (C) of JIMT-1 variants.* *The data are presented as mean ± SD. All analyses were performed in triplicate.*

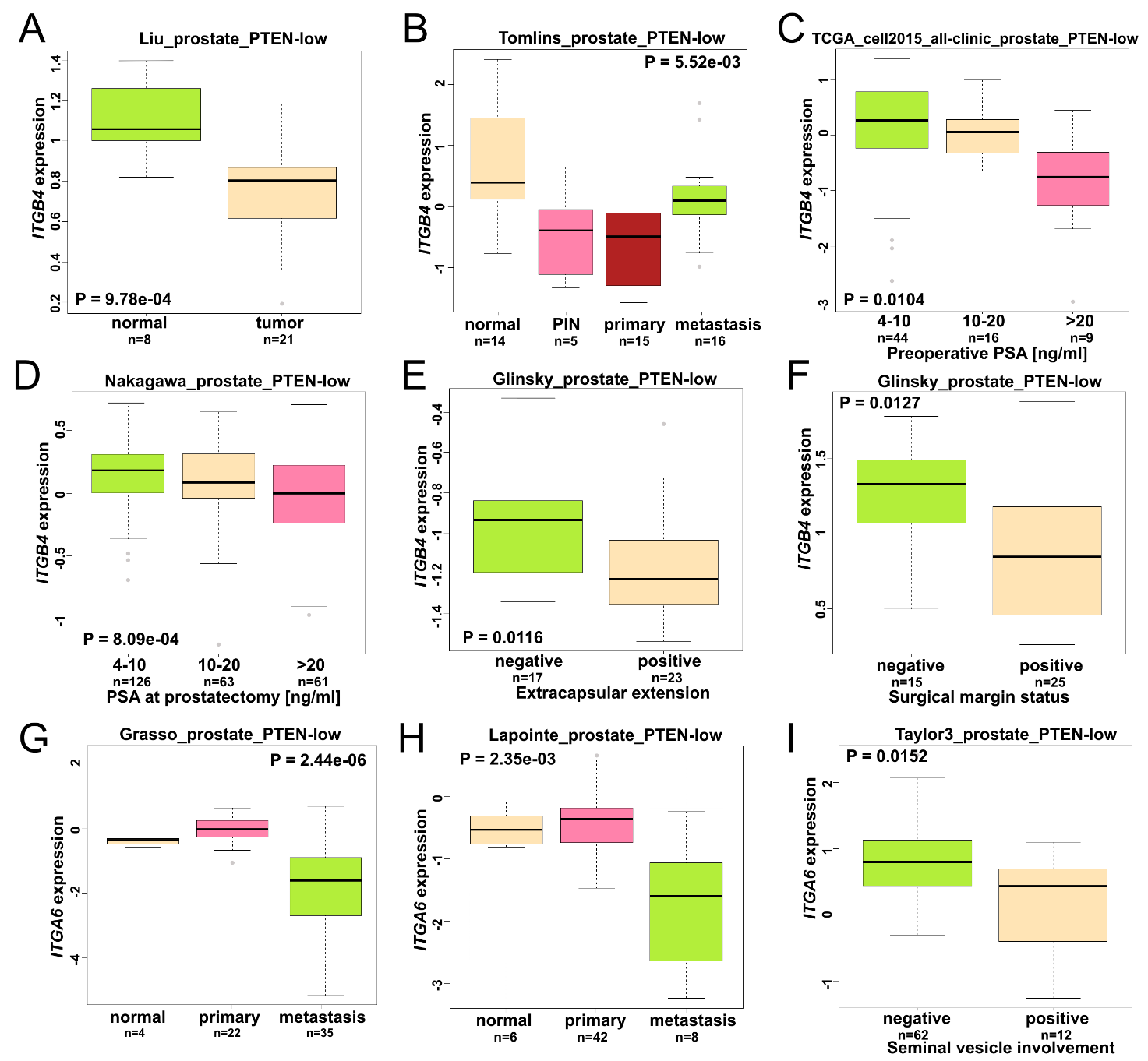

***Figure. S4 Downregulation of ITGB4 or ITGA6 correlates with tumor aggressiveness in PCa patients with low PTEN expression.*** *Lower ITGB4 level correlates with prostate tumor (A), PCa progression (B), PSA levels (C-D), extracapsular extension (E) and surgical margin status (F). Lower ITGA6 level correlates with increased PCa metastasis (G-H) and seminal vesicle involvement (I).* *Statistical tests were assessed by Mann-Whitney U test or Kruskal-Wallis H test depending on the number of comparison groups.*

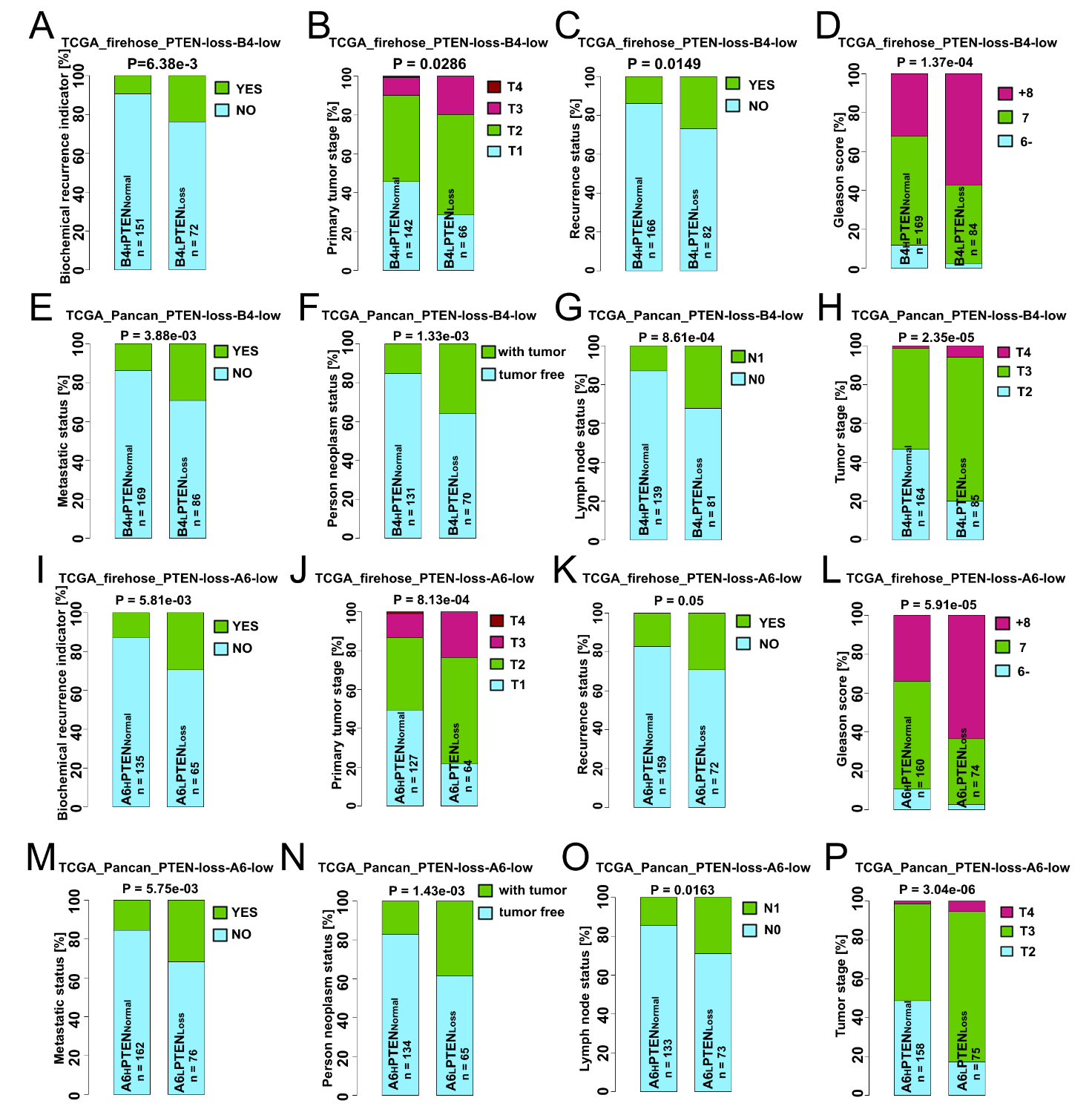

***Figure S5. Simultaneous loss of PTEN and ITGB4/ITGA6 reduction increases tumor aggressiveness in PCa patients.*** *Lower ITGB4 level correlates with biochemical PCa recurrence indicator (A), increased primary tumor stage (B) recurrence status (C), higher Gleason score (D), increased PCa metastasis (E), neoplasm status (F), lymph node metastasis (G) and tumor stage (H). Lower ITGA6 level correlates with biochemical PCa recurrence indicator (I), increased primary tumor stage (J) PCa recurrence status (K), higher Gleason score (L), increased metastasis (M), neoplasm status (N), lymph node metastasis (O) and tumor stage (P).* *The analysis was conducted by Fisher’s exact test, P value < 0.05 was considered as statistically significant.*

***Table S1 List of antibodies used in this study****.*

|  | **Name** | **Company** | **Cat. No** | **Dilution for WB** | **Dilution for ICC/IHC** |
| --- | --- | --- | --- | --- | --- |
| 1 | anti-Akt | Cell Signalling Technology | 9272 | 1:1000 |  |
| 2 | anti-CD151 | gift from Christopher S. Stipp | - |  | 1:100 |
| 3 | anti-cleaved caspase 3 | Cell Signalling Technology | 9661 | 1:750 |  |
| 4 | anti-cleaved PARP | Cell Signalling Technology | 5625 | 1:1000 |  |
| 5 | anti-EGFR | Cell Signalling Technology | 4267 | 1:1000 |  |
| 6 | anti-FAK | BD Transduction Laboratories | 610088 | 1.1000 |  |
| 7 | anti-keratin 8, phospho S431 | Abcam | ab109452 |  | 1:100 |
| 8 | anti-laminin 322 | gift from Karl Tryggvason | - |  | 1:100 |
| 9 | anti-MAPK | Cell Signalling Technology | 9212 | 1:750 |  |
| 10 | anti-pAkt S473 | Cell Signalling Technology | 9271 | 1:750 |  |
| 11 | anti-paxillin | BD Transduction Laboratories | 610051 | 1:1000 |  |
| 12 | anti-pEGFR Y1068 | Cell Signalling Technology | 2234 | 1:750 |  |
| 13 | anti-pERK1/2 T202/Y204 | Cell Signalling Technology | 4370 | 1:750 |  |
| 14 | anti-pFAK Y397 | ECM Bioscience | FM1211 | 1:1000 |  |
| 15 | anti-PI3K | Cell Signalling Technology | 4292 | 1:1000 |  |
| 16 | anti-plectin | Sigma | HPA029906 | 1:1000 | 1:100 |
| 17 | anti-plectin | Epitomics | 1399-1 |  | 1:100 |
| 18 | anti-pMAPK T180/Y182 | Cell Signalling Technology | 4511 | 1:1000 |  |
| 19 | anti-pPI3K Y199 | Cell Signalling Technology | 4228 | 1:750 |  |
| 20 | anti-pSrc Y416 | Cell Signalling Technology | 6943 | 1:1000 |  |
| 21 | anti-PTEN | Santa Cruz Biotechnology | sc7974 | 1:750 |  |
| 22 | anti-PTEN | Cell Signalling Technology | 9559 |  | 1:100 |
| 23 | anti-Src | Cell Signalling Technology | 2109 | 1:1000 |  |
| 24 | anti-vimentin | Dako | M0725 | 1:750 |  |
| 25 | anti-vinculin | NovusBio | NB600-1293 | 1:750 |  |
| 26 | anti-α6 integrin | Sigma | HPA012696 | 1:2000 | 1:100 |
| 27 | anti-α6 integrin | BD Transduction Laboratories | 555734 |  | 1:100 |
| 28 | anti-β1 integrin | gift from Karl Matlin | - |  |  |
| 29 | anti-β3 integrin | BioRAD | MCA2263 |  |  |
| 30 | anti-β4 integrin | Abcam | ab29042 |  | 1:100 |
| 31 | anti-β4 integrin | Santa Cruz Biotechnology | sc-9090 | 1:1000 |  |
| 32 | anti-β-tubulin | Sigma | T4026 | 1:5000 |  |

***Table S2 List of qPCR primers.***

| **Gene** | **Sequence 5’-3’** | **Predicted product size [bp]** |
| --- | --- | --- |
| *CENPA* | F: TCCTTAGGCGCTTCCTCCC  R: CAAGAGGTGTGTGCTCTTCTGA | 90 |
| *DIAPH3* | F: GCTTTTAAGTCTCAGTTTGGTGCC  R: GACCACTGAATGGCATCCGC | 190 |
| *MAD2L1* | F: GCCGAAATCGTGGCCGAG  R: GTTACAAGCAAGGTGAGTCCGT | 119 |
| *INCENP* | F: GAGCTGATGCCCAAAACACCT  R: TGCGGGATAACCTTCTCCTGAT | 106 |
| *CCND1* | F: CCTGTGCTGCGAAGTGGAAA  R: GAAGACCTCCTCCTCGCACT | 220 |
| *GAPDH* | F: AACAGCGACACCCATCCTC  R: CATACCAGGAAATGAGCTTGACAA | 85 |
